# A p53-dependent translational program directs tissue-selective phenotypes in a model of ribosomopathies

**DOI:** 10.1101/2020.06.24.167940

**Authors:** Gerald C. Tiu, Craig H. Kerr, Craig M. Forester, Pallavi S. Krishnarao, Hannah D. Rosenblatt, Nitin Raj, Olena Zhulyn, Margot E. Bowen, Leila Shokat, Laura D. Attardi, Davide Ruggero, Maria Barna

**Author notes:** These authors contributed equally to this work (Gerald C. Tiu, Craig H. Kerr).

## Abstract

In ribosomopathies, perturbed expression of ribosome components leads to tissue-specific phenotypes, such as limb and craniofacial defects as well as bone marrow failure. What accounts for such tissue-selective manifestations as a result of mutations in the ribosome, a ubiquitous cellular machine, has remained a mystery. Combining comprehensive mouse genetics and *in vivo* ribosome profiling, we observe limb patterning phenotypes in ribosomal protein (RP) haploinsufficient embryos and uncover corresponding selective translational changes of transcripts controlling limb development. Surprisingly, both loss of p53, which is activated by RP haploinsufficiency, and augmented protein synthesis rescue these phenotypes. These findings are reconciled by the unexpected identification that p53 functions as a master regulator of protein synthesis through transcriptional activation of 4E-BP1. 4E-BP1, a key regulator of translation, in turn, facilitates selective changes in the translatome downstream of p53 and thereby explains, at least in part, how RP haploinsufficiency elicits specificity to gene expression. These results provide an integrative model to explain how *in vivo* tissue-specific phenotypes emerge from a mutation in a ribosome component.

## INTRODUCTION

Ribosomopathies are a class of diseases characterized by mutations in ribosome components or key ribosome biogenesis factors that produce unexpected tissue-specific phenotypes. For example, Diamond Blackfan Anemia (DBA), which arises from haploinsufficiency of ∼20 different essential ribosomal proteins, converges upon a common set of poorly understood congenital birth defects such as craniofacial, digit, and limb abnormalities, as well as selective impairment of erythroid differentiation. The observation that perturbations in a number of different core ribosomal proteins converge on a common set of disease phenotypes suggests that general ribosome dysfunction underlies selective disease manifestations in DBA (Clinton and Gazda, 1993; Yelick and Trainor, 2015). While the disease phenotypes in DBA have been the topic of intense study, the mechanisms underlying them, particularly congenital birth defects, are still poorly understood.

Many studies suggest that *in vivo* phenotypes underlying ribosomopathies can arise from the activation of the stress-induced transcription factor p53. In particular, mutations in ribosomal proteins are thought to lead to p53 stabilization and activation through a mechanism in which nucleolar stress and defective ribosome biogenesis result in inhibition of the major p53 negative regulator, Mdm2 (Deisenroth and Zhang, 2010; Dutt et al., 2011; Morgado-Palacin et al., 2015; Narla and Ebert, 2010). Stabilization of p53 is thought to contribute, at least in part, to ribosomopathies through transcriptional activation of genes canonically involved in regulating cell proliferation and apoptosis (Deisenroth and Zhang, 2010). The role of p53 in the generation of these phenotypes is underscored when examining RP haploinsufficiency within an *in vivo* context. Studies in both mice and zebrafish models of DBA have demonstrated that ribosomal protein haploinsufficiency can lead to tissue-specific defects *in vivo*, which can be rescued upon loss of p53 (Barlow et al., 2010; Danilova et al., 2008; McGowan and Mason, 2011; McGowan et al., 2008; Sulic et al., 2005). While these findings highlight the role of p53 activation in the development of disease phenotypes, how p53-mediated global effects on cells, such as cell cycle arrest and apoptosis, can lead to tissue-specific developmental phenotypes remains a paradox.

Alternatively, it has been suggested that mutations in ribosomal proteins may lead to translational dysregulation independent of p53 activation (Dalla Venezia et al., 2019; Khajuria et al., 2018). Despite this, changes to translation have been largely unexplored within *in vivo* developmental contexts associated with RP haploinsufficiency. A recent *ex vivo* cell culture study identified translational defects in hematopoietic cells upon knockdown of RPs mutated in DBA, and suggests that ribosome levels may play a role in this dysfunction (Khajuria et al., 2018). However, it is perplexing how this hypothesis accounts for complex *in vivo* phenotypes and how it can be reconciled with the known role of p53 in the etiology of ribosomopathy phenotypes.

Taken together, it remains a mystery what roles p53-dependent and - independent mechanisms play in the development of the *in vivo* tissue-specific phenotypes associated with loss of function of a core ribosome component. Here, we genetically inactivate one allele of the core ribosomal protein, *Rps6*, in the developing mouse limb bud as a model system for investigating the complex phenotypes associated with ribosomopathies. Strikingly, we find that *Rps6* haploinsufficiency leads to selective limb phenotypes reminiscent of those observed in DBA. By combining comprehensive mouse genetics and *in vivo* ribosome profiling, we demonstrate that translational dysfunction plays a critical role in development of tissue-specific phenotypes upon *Rps6* haploinsufficiency. Moreover, our ribosome profiling uncovered hundreds of transcripts which undergo differential translational regulation upon *Rps6* haploinsufficiency. The majority of these transcript-specific translational changes are unexpectedly rescued upon loss of p53, indicating that translational control and p53 activation are in fact coupled. To this end, we demonstrate that p53 in addition to its bona fide role in transcriptional regulation, is also a master regulator of protein synthesis. This function is, at least in part, mediated through p53-dependent induction of a key translational regulator, 4E-BP1 (eukaryotic initiation factor 4E-binding protein 1), which is known to have a role in translational control of selective mRNAs (Thoreen et al., 2012; Truitt et al., 2015). Together this work provides an integrative model wherein mutations in core components of the ribosome activate p53 to directly lead to transcript-specific changes in cap-dependent translation by modulating 4E-BP1 expression. Overall, these findings provide key insight into how mutations of core ribosome components underlie tissue-specific phenotypes in ribosomopathies.

## RESULTS

### Ribosomal protein haploinsufficiency leads to selective developmental phenotypes

To systematically delineate how haploinsufficiency of an essential RP leads to tissue-specific developmental phenotypes, we perturbed expression of the small subunit RP, Rps6 (Panić et al., 2006; Volarevic et al., 2000), in the developing mouse embryo. Rps6 loss of function has served as a model to study ribosomopathies, as conditional *Rps6* hemizygosity within the hematopoietic compartment leads to erythroid phenotypes similar to that of other RPs found mutated in ribosomopathies, such as in DBA (McGowan et al., 2011). However, *Rps6* haploinsufficiency at the whole-organismal level is embryonic lethal (Panić et al., 2006). Therefore, to investigate the outstanding question of how congenital birth defects arise in an *in vivo* context due to RP haploinsufficiency, without affecting organismal viability, we turned to the developing limb bud. Limb development is a well-established model system for studying tissue patterning and where defects emerge in ribosomopathies (Gazda et al., 2008). Importantly, patients diagnosed with DBA have been characterized with limb defects including hypoplastic digit 1 and radial abnormalities (Gazda et al., 2008; Hurst et al., 1991).

During development, limbs arise from a small bud of mesenchymal cells that includes chondrocyte progenitors, which serve as precursors to all the skeletal elements of the limb covered by a surface ectoderm. The apical ectodermal ridge (AER) is a morphologically distinct ectoderm surrounding the distal tip of the limb bud that serves as an important signaling center that promotes limb outgrowth and survival of the underlying mesenchyme (Zeller et al., 2009) (Figure 1A). The basic vertebrate limb skeleton derives from a cartilaginous model: a single proximal long bone within the stylopod segment (humerus; femur), followed by two long bones within the zeugopod segment (radius, ulna; tibia, fibula) and then the distal autopod segment comprising of wrist or ankle, and digits (Figure 1A). Using two distinct and well-established Cre lines driven by the *Prx1* promoter/enhancer (Logan et al., 2002) or the *Msx2* promoter (Sun et al., 2000), we conditionally deleted one allele of *Rps6* in either the underlying limb mesenchyme or AER, reflecting the two major cell lineages that either give rise to the limb cartilage template or control limb outgrowth and skeletal patterning, respectively.

**Figure 1.**
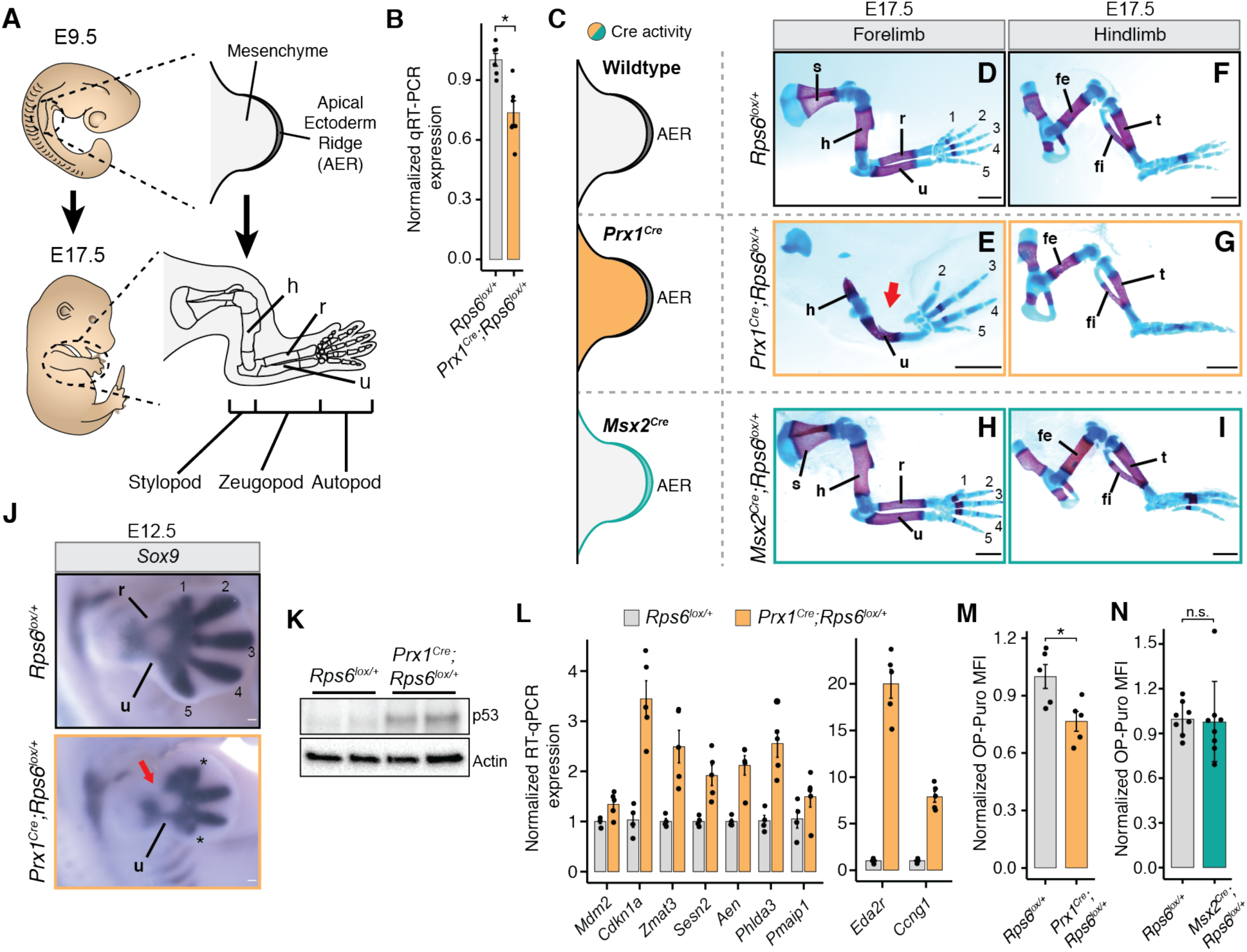
*Rps6* haploinsufficiency in the developing limb bud mesenchyme leads to selective patterning defects marked by activation of p53 and a reduction in global protein synthesis. (**A**) Overview of forelimb development from embryonic day 9.5 (E9.5) to E17.5. AER, apical ectodermal ridge; h, humerus; r, radius; u, ulna. (**B**) RT-qPCR of *Rps6* mRNA in forelimbs from E10.5 embryos. Expression normalized to the geometric mean of housekeeping genes *Actb, Tbp, Ubb, Exo5, Pkm2,* and *Nadk2* then to the mean of wildtype values. n = 6. (**C**) Diagram of Cre recombinase activity distribution in limb bud mesenchyme (*Prx1^Cre^*; orange), or in the overlying AER (*Msx2^Cre^*; green). (**D-I**), Representative images of E17.5 forelimbs and hindlimbs from wildtype (top row), *Prx1^Cre^;Rps6^lox/+^* (middle row), and *Msx2^Cre^;Rps6^lox/+^* (bottom row). Bone, red; Cartilage, blue. Numbers indicate digits. Arrow indicates absence of radius. Scale bars, 1 mm. (**J**) *Sox9 in situ* hybridization of E12.5 forelimbs. Numbers indicate mesenchymal condensations leading to digit formation. *Impaired digit development. Arrow indicates absence of radius. Scale bars, 0.1 mm. (**K**) Representative Western blot of p53 in E10.5 forelimbs from 2 separate embryos. (**L**) RT-qPCR of p53 target genes in E10.5 forelimbs. Expression was normalized to *Actb* and to the mean of wildtype values. *Eda2r* and *Ccng1* were plotted separately for clarity given higher expression values relative to other p53 target genes. *n* = 4 embryos (*Rps6^lox/+^*), *n* = 5 embryos (*Prx1^Cre^;Rps6^lox/+^*). (**M**) OP-Puro MFI of cells dissociated from E10.5 forelimbs of *Prx1^Cre^;Rps6^lox/+^* mice normalized to wildtype. *n* = 5 embryos; MFI = median fluorescence intensity. (**N**) OP-Puro MFI of ectodermal cells after separation of the ectoderm from E11.5 forelimbs of *Msx2^Cre^;Rps6^lox/+^* mice normalized to wildtype. *n* = 8 embryos; MFI = median fluorescence intensity. For all bar plots, error bars = SEM, * *P* < 0.05, ** *P* < 0.01, *** *P* < 0.001, **** *P* < 0.0001, two-tailed t-test, unequal variance.

Within the forelimb of *Prx1^Cre^;Rps6^lox/+^* embryos, a significant decrease of *Rps6* mRNA and protein was detected as early as day 10.5 of embryonic development (E10.5) where Cre recombinase was broadly activated throughout the limb bud mesenchyme (Figure 1B-C; Figure S1A; Figure S7A-B) (Logan et al., 2002). At E17.5, *Prx1^Cre^;Rps6^lox/+^* embryos demonstrated striking selective underdevelopment (hypoplasia) or loss (aplasia) of the anterior zeugopod element (the radius) compared to the ulna (28/28 limbs) in addition to scapular hypoplasia (28/28 limbs); hypoplasia/absence of the proximal humerus (28/28 limbs); and selective hypoplasia/absence of digits 1/2 (28/28 limbs), digit 4 (26/28 limbs), and digit 5 (28/28 limbs) marked by reduced phalangeal numbers, with digits 1 and 5 exhibiting the most severe phenotypes (Figure 1D-E; Figure S1C-D; Table S1). Quantifying the ratio of radial ossification length to that of the ulna confirmed significant and penetrant radial aplasia in the *Prx1^Cre^;Rps6^lox/+^* embryos compared to control (Figure S1B). Interestingly, the *Prx1^Cre^;Rps6^lox/+^* phenotype is reminiscent of those found in DBA patients, who can present with limb patterning defects including hypoplastic digit 1 and selective radial aplasia, as mentioned above (Gazda et al., 2008; Hurst et al., 1991). The hindlimbs, on the contrary, displayed a much milder phenotype in *Prx1^Cre^;Rps6^lox/+^* embryos consisting of a hypoplastic digit 1 and a smaller, medially displaced patella (Figure 1F-G; Figure S1E-F, G-J). Surprisingly, we did not observe any phenotype upon *Rps6* haploinsufficiency in the AER both within the forelimb and hindlimb (see *Msx2^Cre^;Rps6^lox/+^*; Figure 1H-I; Figure S1K-N). In comparison to the selective skeletal phenotypes observed in the *Prx1^Cre^;Rps6^lox/+^* embryos, the lack of phenotype upon *Rps6* haploinsufficiency in the AER in the *Msx2^Cre^;Rps6^lox/+^* embryos further suggests tissue and context specific effects of RP haploinsufficiency.

### Both p53 activation and decreased protein synthesis coincide with limb patterning defects

The selective radial and digit phenotypes in *Prx1^Cre^;Rps6^lox/+^* embryos were observed as early as E12.5 by *in situ* hybridization of *Sox9*, the earliest marker of mesenchymal condensations, which prefigure cartilage elements, suggesting that these phenotypes result from defects in patterning prior to cartilage formation (Figure 1J). We hypothesized that the observed phenotypes may arise from either p53 activation or perturbed protein synthesis. Thus, we first asked whether p53 is activated in E10.5 *Prx1^Cre^;Rps6^lox/+^* limb buds, approximately one day after Cre-mediated *Rps6* recombination in the limb mesenchyme. Indeed, we observed spatially homogeneous p53 activation and transcriptional upregulation of known p53 target genes (Figure 1K-L; Figure S2A) (Bowen et al., 2019; Brady et al., 2011). This was accompanied by increased apoptosis and mildly decreased cell proliferation that were not spatially restricted within the limb bud mesenchyme (Figure S2B-E). We also observed p53 activation in the ectoderm of *Msx2^Cre^;Rps6^lox/+^* embryos (Figure S3). Next, we measured changes in global protein synthesis per cell by quantifying incorporation of O-propargyl-puromycin (OP-Puro) into nascent peptides of freshly dissociated limb mesenchymal cells at stage E10.5 (Figure S4) (Signer et al., 2014). We observed reduced OP-Puro incorporation in *Prx1^Cre^;Rps6^lox/+^* versus control, indicating that protein synthesis is impaired in the developing limb mesenchyme upon Rps6 reduction (Figure 1M). By contrast, we did not observe a decrease in OP-Puro incorporation when we assayed the cells derived from separated ectoderm layer of *Msx2^Cre^;Rps6^lox/+^* limb buds (Figure 1N), despite p53 activation and decreased *Rps6* mRNA levels (Figure S3). Taken together, these data suggest that both p53 activation and defects in global protein synthesis may be associated with the observed limb patterning phenotypes upon *Rps6* haploinsufficiency. Moreover, as *Rps6* haploinsufficiency in the mesenchyme leads to phenotype and a decrease in protein synthesis whereas haploinsufficiency in the AER does not, our findings also suggest that diminished protein synthesis may be an important driver of tissue selective phenotypes observed upon RP haploinsufficiency.

### mTORC1 activation or loss of p53 rescue limb patterning defects

As both p53-activation and translational dysfunction accompany the *Prx1^Cre^;Rps6^lox/+^* limb phenotypes, we next turned to *in vivo* genetics to determine the contributions of decreased protein synthesis and/or p53 activation on the *Prx1^Cre^;Rps6^lox/+^* limb phenotypes (Figure 2A). First, we genetically modulated global protein synthesis using one of the most well-described pathways that impacts translational control, the mTOR pathway (Saxton and Sabatini, 2017). mTOR signaling functions are distributed between at least two distinct mTOR protein complexes: mTORC1 and mTORC2. In particular, one of the major functions of mTORC1 signalling is controlling protein synthesis levels by phosphorylating key translational regulators, which subsequently leads to upregulation of cap-dependent protein synthesis (Figure 2B) (Saxton and Sabatini, 2017; Signer et al., 2014; Zhang et al., 2014). By genetically modulating the activity of mTORC1, and consequently its ability to upregulate translation, we could address whether translation is implicated in the development of *Prx1^Cre^;Rps6^lox/+^* limb phenotypes. To specifically activate mTORC1 in the limb, we conditionally inactivated the tuberous sclerosis complex (TSC), which specifically represses mTORC1 (Garami et al., 2003; Zhang et al., 2014), via genetic deletion of the TSC component *Tsc2*. Remarkably, deletion of *Tsc2* in developing limbs (Figure S5A) rescued the *Rps6* haploinsufficiency radius patterning phenotype as exemplified by improved radial development in *Prx1^Cre^;Rps6^lox/+^;Tsc2^lox/lox^* compared to *Prx1^Cre^;Rps6^lox/+^;Tsc2^+/+^* limbs (Figure 2C-F; Figure S5B). This phenotypic rescue in limb patterning is accompanied by a concurrent increase in protein synthesis as measured by OP-Puro incorporation in E10.5 limb buds (Figure 2G). While we only observed a partial rescue of the limb phenotype by *Tsc2* deletion, this may be explained by the incomplete restoration of protein synthesis back to wildtype levels upon loss of *Tsc2* (Figure 2G). In addition, loss of *Tsc2* did not influence *Rps6* expression nor does it abrogate the p53 activation induced by *Rps6* haploinsufficiency (Figure 2H; Figure S5A,C-D). Taken together, these results suggest that perturbed translation may contribute to the developmental patterning phenotypes observed upon Rps6 reduction.

**Figure 2.**
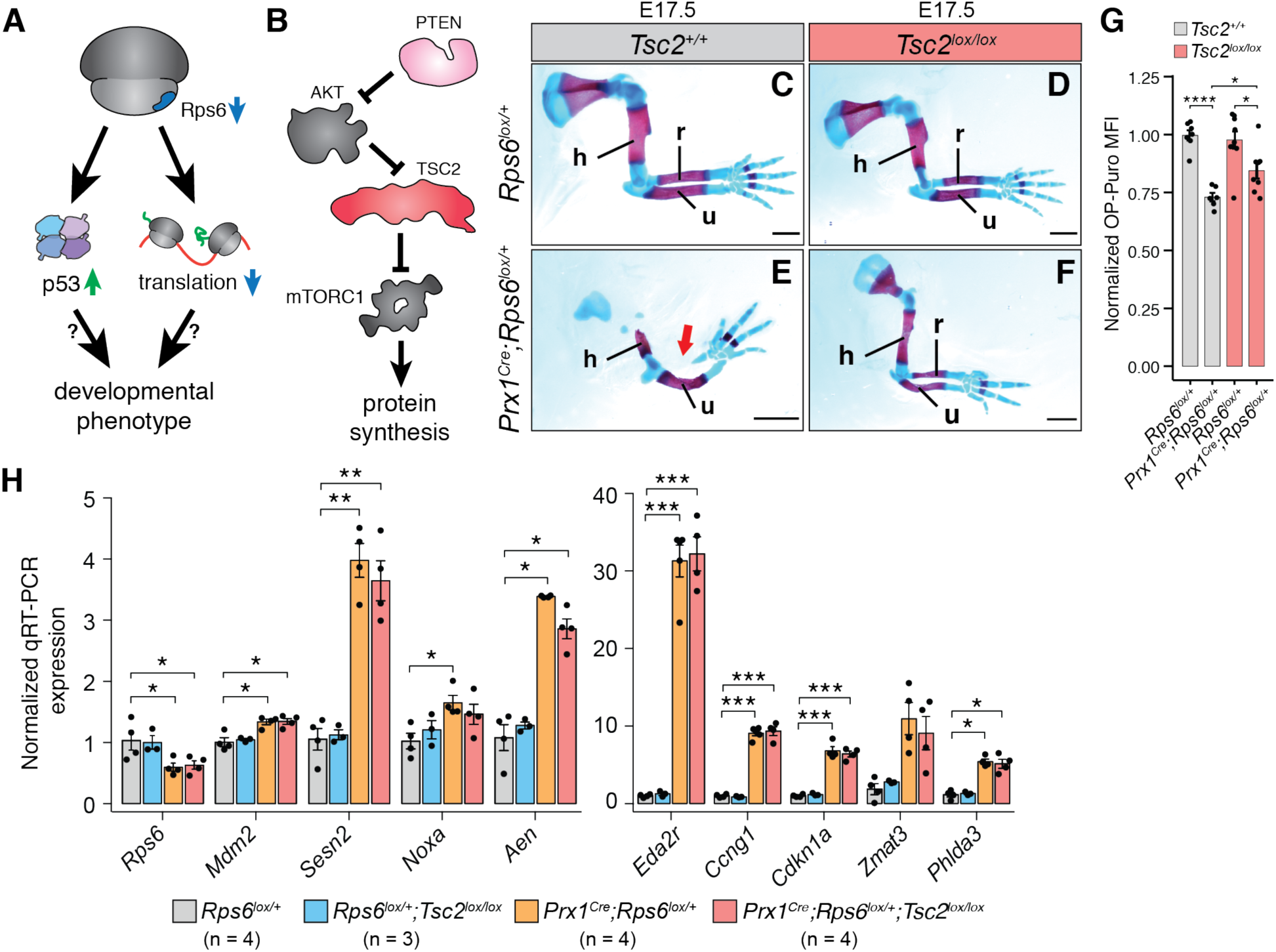
mTORC1 activation with corresponding augmented protein synthesis rescues phenotypes associated with *Rps6* haploinsufficiency. (**A**) Overview of pathways potentially leading to developmental phenotypes upon *Rps6* haploinsufficiency. p53 activation and impaired translation have been proposed as independent models to explain developmental phenotypes in ribosomopathies. In Figure 2, we investigate the contribution of translation dysregulation. (**B**) Schematic of mTORC1 regulation and downstream effects. (**C-F**) Representative E17.5 forelimbs of wildtype (*Rps6^lox/+^*) and *Prx1^Cre^;Rps6^lox/+^* embryos in *Tsc2* wildtype (*Tsc2^+/+^*) or conditional loss of *Tsc2* (*Tsc2^lox/lox^*) background. Arrow indicates absence of radius. Scale bars, 1 mm; h, humerus; r, radius; u, ulna. (**G**) OP-Puro MFI of cells dissociated from E10.5 forelimbs normalized to wildtype. *n* = 7 embryos, (*Rps6^lox/+^;Tsc2^+/+^*); *n* = 6 embryos, (*Prx1^Cre^;Rps6^lox/+^;Tsc2^+/+^*); *n* = 9 embryos, (*Rps6^lox/+^;Tsc2^lox/lox^*); *n* = 8 embryos, (*Prx1^Cre^;Rps6^lox/+^;Tsc2^lox/lox^*). (**H**) Relative expression of p53 target genes as measured by RT-qPCR of E10.5 forelimbs. Expression is normalized to the geometric mean of *Actb* and *Tbp* then to the mean of wildtype values. *n* = 4 embryos for *Rps6^lox/+^, Prx1^Cre^;Rps6^lox/+^, and Prx1^Cre^;Rps6^lox/+^;Tsc2^lox/lox^*, *n* = 3 embryos for *Rps6^lox/+^;Tsc2^lox/lox^*. For all bar plots, error bars = SEM, * *P* < 0.05, ** *P* < 0.01, *** *P* < 0.001, **** *P* < 0.0001, two-tailed t-test, unequal variance.

To formally address the potential contribution of p53 activation to limb patterning defects upon *Rps6* haploinsufficiency in the mesenchyme, we next examined how loss of p53 impacts the limb phenotype (Figure 3). To our surprise, deletion of p53 in the *Prx1^Cre^;Rps6^lox/+^* background also resulted in the rescue of the limb phenotype (Figure 3A-D; Figure S6A-D). Together these findings surprisingly show that either activation of the mTORC1 pathway with associated restoration of global protein synthesis or inactivation of p53 results in a genetic rescue of *Rps6* haploinsufficient patterning phenotypes *in vivo*. These genetic findings raise the possibility that p53 activity and translational control may be unexpectedly coupled in the context of *in vivo* RP haploinsufficiency. In support of this, past studies in cultured cells have suggested that p53 may regulate translation (Kasteri et al., 2018), for example by suppressing mTORC1 activity by activating Sestrin, a negative regulator of mTORC1 (Budanov and Karin, 2008; Loayza-Puch et al., 2013). However, in the context of *Rps6* haploinsufficiency *in vivo*, we do not observe differential mTORC1 activity in *Prx1^Cre^;Rps6^lox/+^* E10.5 limb buds compared to wildtype (Figure S7). Therefore, we explored whether p53 accounts for the alterations in protein synthesis observed upon RP haploinsufficiency through a yet unknown mechanism (Figure 3E). Strikingly, loss of p53 rescues the protein synthesis defect observed upon reduction of Rps6 in E10.5 limb buds as measured by OP-Puro incorporation (Figure 3F). To determine whether p53 activation alone is sufficient to repress global protein synthesis, we next assayed the incorporation of the methionine analog L-Homopropargylglycine (HPG) into nascent peptides in primary mouse embryonic fibroblasts (MEFs) treated with two different p53 small molecule activators. Nutlin-3a specifically blocks Mdm2-mediated p53 destabilization, while doxorubicin (Doxo) induces genotoxic stress (Vassilev et al., 2004; Wang et al., 2004). Nutlin-3a and doxorubicin treatment led to a decrease in global protein synthesis within 8 hours of treatment (Figure 3G). Importantly, we did not observe a significant difference in apoptosis and only a modest decrease in cell proliferation after 8 h of Nutlin-3a treatment (Figure S8A-B). Collectively, these findings unexpectedly demonstrate that activation of p53 is required to repress protein synthesis upon *Rps6* haploinsufficiency and that p53-dependent control of translation occurs through an unknown mechanism independent of mTORC1 pathway activation.

**Figure 3.**
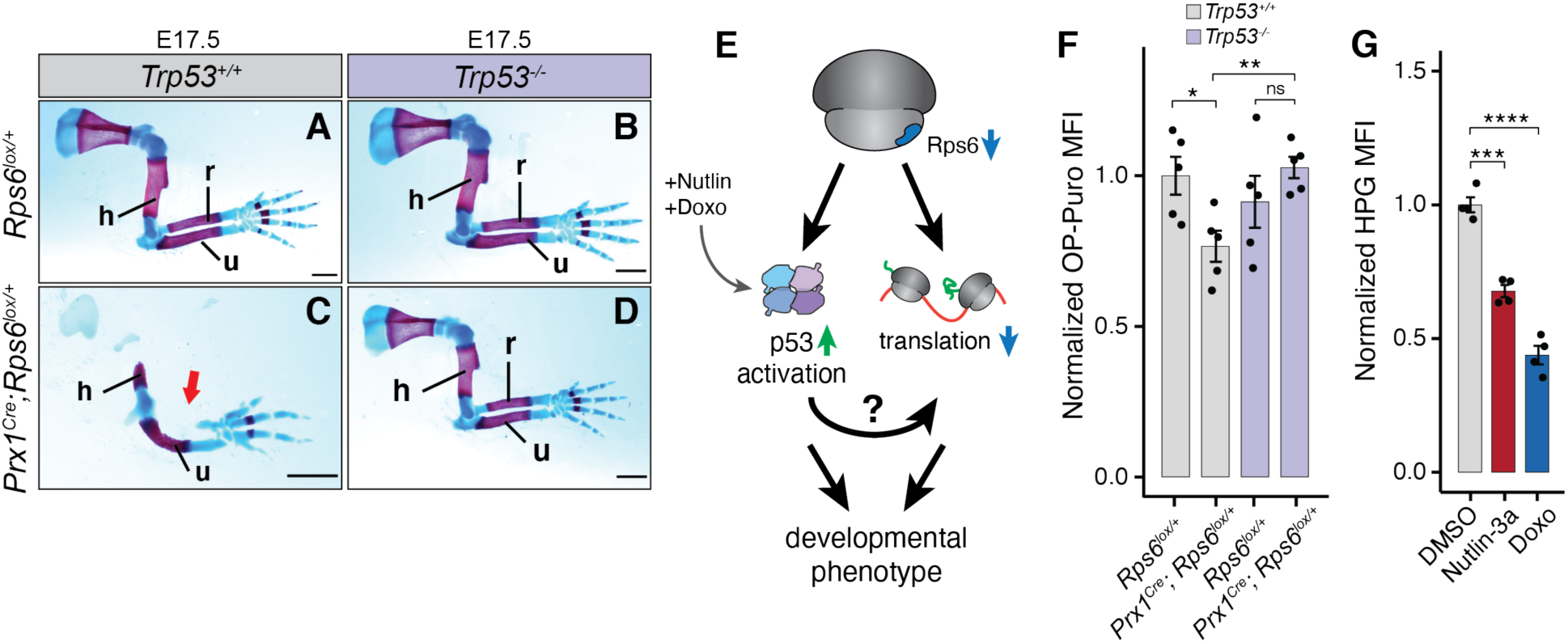
Loss of p53 rescues the *Rps6* haploinsufficiency phenotypes, and p53 activation mediates translational changes upon Rps6 reduction. (**A-D**) Representative E17.5 forelimbs of wildtype and *Prx1^Cre^;Rps6^lox/+^* embryos in a *Trp53* wildtype (*Trp53^+/+^*) and *Trp53* null (*Trp53^-/-^*) background. Arrow indicates absence of radius. Scale bars, 1 mm; h, humerus; r, radius; u, ulna. (**E**) Potential pathways for p53-dependent translational control upon *Rps6* haploinsufficiency. (**F**) OP-Puro MFI of cells dissociated from E10.5 forelimbs normalized to mean of wildtype (*Rps6^lox/+^*). *n* = 5 embryos. (**G**) HPG MFI of mouse embryonic fibroblasts treated with Nutlin-3a or doxorubicin normalized to mean of DMSO treated control. 8 h treatment, *n* = 4. For all bar plots, error bars = SEM, * *P* < 0.05, ** *P* < 0.01, *** *P* < 0.001, **** *P* < 0.0001, two-tailed t-test, unequal variance.

### p53 drives translational changes upon Rps6 haploinsufficiency

Our results suggest that a p53-dependent translational program may contribute to the limb patterning defects observed upon RP haploinsufficiency. We therefore next asked what transcripts may be selectively perturbed upon RP haploinsufficiency in a p53-dependent and -independent manner. To address this question, we performed genome-wide ribosome profiling directly on E10.5 *Rps6* haploinsufficient limb buds in a *Trp53^+/+^* or *Trp53^-/-^* background (Figure 4A-B, Figure S9, Figure S10) (Ingolia et al., 2011). Although standard ribosome profiling cannot detect alterations in global protein synthesis, this technique can pinpoint relative changes in the translation efficiency (TE) of specific transcripts (McGlincy and Ingolia, 2017). We observed a large degree of translational changes upon Rps6 reduction (282 transcripts TE decreased, 138 transcripts TE increased) of which most changes are rescued by loss of p53 (Figure 4A-B, Figure S10A,C, Table S3-4). Interestingly, the TE changes of a smaller subset of differentially translated mRNAs, which included p53 regulators and cytoplasmic RPs (see below), were not dependent on p53 (64 transcripts TE increased, 8 transcripts TE decreased) (Figure 4A-B, Figure S10B,D, Table S4).

**Figure 4.**
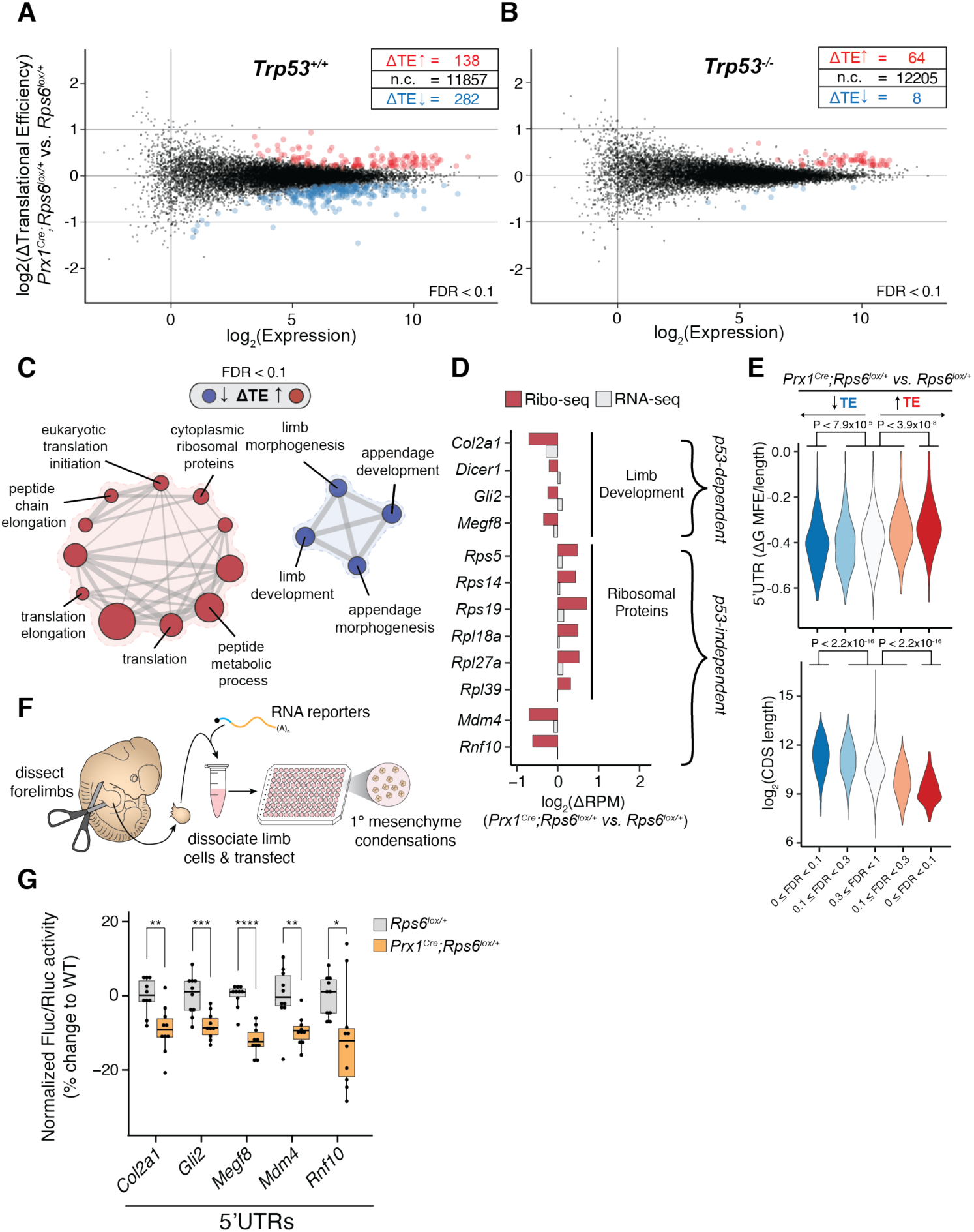
*In vivo* ribosome profiling reveals the landscape of p53-dependent and -independent translation changes upon *Rps6* haploinsufficiency. (**A-B**) MA plot of change in translational efficiency (ΔTE) in E10.5 *Prx1^Cre^;Rps6^lox/+^* vs. *Rps6^lox/+^* embryonic forelimbs in *Trp53^+/+^* background (**A**) and *Trp53^-/-^* background (**B**). red, ΔTE > 0; blue, ΔTE < 0; *n* = 3 biological replicates (2 embryos each); FDR < 0.1. (**C**) Gene sets enriched for transcripts changing in TE in E10.5 *Prx1^Cre^;Rps6^lox/+^* vs. *Rps6^lox/+^* forelimbs. Analysis performed by CAMERA, transcripts filtered for expressed genes; node size, gene set size; edge size, gene set overlap; red, ΔTE > 0; blue, ΔTE < 0; FDR < 0.1. (**D**) Relative change in ribosome footprints for select high-confidence transcripts (red; FDR < 0.1). Relative changes in mRNA expression shown in grey. (**E**) Violin plots quantifying ΔTE relative to 5’UTR structuredness as predicted by UCSC genome browser foldUtr5 normalized to UTR length (ΔG MFE / 5’UTR length) and ΔTE relative to CDS length. Transcripts are stratified by direction of ΔTE (blue, down; red, up) and FDR (0 ≤ FDR < 0.1, 0.1 ≤ FDR < 0.3, 0.3 ≤ FDR ≤ 1); Mann-Whitney U test. (**F**) Workflow of the primary limb micromass assay. (**G**) Firefly/Renilla luciferase activity from transfection of RNA reporters in E11.5 limb micromass assays normalized to the geometric mean of control *Pkm* and *Cnot10* 5’UTR activities and wildtype (*Rps6^lox/+^*). *Pkm* and *Cnot10* were chosen as controls given that they did not change in translation efficiency upon *Rps6* haploinsufficiency in the ribosome profiling experiment (Table S3). For box-plots, center line, median; box limits, first and third quartiles; whiskers, 1.5x interquartile range; points, outliers. * *P* < 0.05, ** *P* < 0.01, *** *P* < 0.001, **** *P* < 0.0001, two-tailed t-test, unequal variance.

Gene set enrichment analysis revealed that translationally repressed gene sets include those involved in limb development (Figure 4C, Table S5), which correlates with the observed phenotypes. Hence, the selective changes in translational control may account for the specificity in the limb patterning defects observed. For example, high confidence transcripts in these gene sets whose decrease in TE is primarily due to reduction in ribosome footprints and not to changes in transcript levels included *Col2a1*, *Dicer1*, *Gli2*, and *Megf8* (Figure 4D, see Materials and Methods for selection criteria). *Col2a1* encodes for collagen type II alpha 1 chain, a component of the cartilage that lays the framework for skeletal development of most bones (Lee et al., 1989). *Dicer1*, which encodes an RNase essential for miRNA maturation, is essential for proper limb maturation and survival of mesenchymal cells (Harfe et al., 2005). Furthermore, loss of *Gli2*, a component of the Sonic Hedgehog pathway, leads to shortened limbs with particular shortening of the radial bone (Mo et al., 1997), while mutations in *Megf8* lead to limb developmental defects in humans (Twigg et al., 2012). In addition, translational repression of these select transcripts involved in limb development is p53-dependent, consistent with the requirement of p53 for the development of limb phenotypes upon *Rps6* haploinsufficiency (Figure 4D; Figure S10A,C).

Translationally upregulated mRNAs were enriched in gene sets that included those containing cytoplasmic RPs (Figure 4C-D; Figure S10B,D, Table S5). In contrast to the translationally repressed limb development transcripts that are p53-dependent, RP translational upregulation is p53-independent (50 RPs out of 64 p53-independent upregulated transcripts, Figure 4D, Figure S10B,D, Table S4). Our *in vivo* results may thus represent a possible translational feedback mechanism to restore RP homeostasis when core RP expression is perturbed. In fact, translation of *Rps6* itself was upregulated (Table S3-4), suggesting that the phenotype may in fact be more severe with a further decrease in Rps6 protein levels had it not been for this feedback mechanism. Upregulation of RP translation was particularly unexpected given that *ex vivo* studies demonstrated a marked decrease in RP translation after RP depletion (Khajuria et al., 2018), suggesting that results from *in vivo* RP haploinsufficiency models may differ profoundly from RP knockdown in cultured cells. We also observed p53-independent translational downregulation of the p53 negative regulator *Mdm4* and of the RING finger-containing protein, *Rnf10* (Figure 4D, Figure S10B,D, Table S4). Mdm4, in particular, cooperates with Mdm2 to suppress p53 activity (Marine et al., 2007). Thus, translational suppression of *Mdm4* may serve as a post-transcriptional quality control sensor to activate p53 in the context of RP haploinsufficiency.

We next validated our ribosome profiling results using sucrose gradient polysome fractionation and subsequent RT-qPCR on select, differentially translated transcripts from E10.5 *Prx1^Cre^;Rps6^lox/+^* and *Rps6^lox/+^* limb buds (Figure S11). Upon *Rps6* haploinsufficiency, we observed a shift to lighter polysomes for transcripts whose TE decreased in the ribosome profiling (e.g., *Col2a1, Dicer1, Gli2, Megf8, Mdm4,* and *Rnf10*) and observed a shift towards heavier polysomes for transcripts whose TE increased in the ribosome profiling (e.g., RPs) (Figure S11B-C). Taken together, these data demonstrate that complex translational remodeling occurs upon *Rps6* haploinsufficiency that consists of a p53-dependent program associated with transcripts that correlate with phenotypic changes and a smaller p53-independent program that may mediate homeostatic responses to RP perturbations.

Next, we investigated what features may regulate differential translational sensitivity of transcripts to *Rps6* haploinsufficiency. Previous studies have demonstrated that *cis*-features such as ORF length and 5’UTR structuredness contribute to translational control (Weinberg et al., 2016). Indeed, we find that translationally repressed transcripts upon Rps6 reduction have longer ORFs compared to the pool of unchanging transcripts whereas transcripts that are upregulated in translation have shorter ORFs (Figure 4E, Figure S12). Additionally, repressed transcripts have more structured 5’UTRs whereas upregulated mRNAs tend to have less structured 5’UTRs (Figure 4E). Notably, the p53-dependent, translationally repressed limb developmental genes *Col2a1*, *Dicer1*, *Gli2*, and *Megf8* have long ORF lengths and highly structured predicted 5’UTRs (Table S6).

Given these observations, we asked whether the 5’UTRs of differentially translated mRNAs contribute to differential translation in response to *Rps6* haploinsufficiency. To this end, we cloned candidate 5’UTRs into luciferase reporter constructs and transfected *in vitro* transcribed Firefly luciferase mRNA along with control Renilla luciferase mRNA into primary limb micromass cultures derived from *Rps6^lox/+^* or *Prx1^Cre^;Rps6^lox/+^* limb buds (Figure 4F). Primary limb micromass cultures are a well-established model system to study limb mesenchyme and cartilage formation that mimics many of the *in vivo* steps of cellular differentiation (Barna and Niswander, 2007). In comparison to two control 5’UTRs taken from transcripts whose translation was unaffected by *Rps6* haploinsufficiency (*Pkm* and *Cnot10*), we observed reduced reporter activity driven by the 5’UTRs of translationally repressed transcripts including *Col2a1*, *Gli2, Megf8, Mdm4*, and *Rnf10* in *Prx1^Cre^;Rps6^lox/+^* limb micromass cultures (Figure 4G). These results are in agreement with our ribosome profiling and polysome RT-qPCR data and suggest that 5’UTRs may drive the translational regulation of these genes. Overall, these findings demonstrate that 5’UTRs are important *cis*-regulatory elements that may contribute, at least in part, to differential translational sensitivity to RP haploinsufficiency.

### p53 modulates translational control by upregulating 4E-BP1 expression

We next sought to determine mechanistically how RP haploinsufficiency leads to p53-dependent translational remodeling (Figure 5A). Thus, we first investigated whether these translational changes depend on the transcriptional transactivation activity of p53. Indeed, in MEFs expressing the p53 transactivation dead mutant *Trp53^LSL-25,26,53,54/LSL-25,26,53,54^ (Trp53^QM/QM^)* (Brady et al., 2011) we did not observe a reduction in protein synthesis (Figure 5B; Figure S13A) after treatment with Nutlin-3a or doxorubicin, suggesting that p53-dependent translational repression requires p53 transactivation activity. We therefore asked what p53 transcriptional targets beyond those previously described may be responsible for these translational changes. Strikingly, we found that transcription of *Eif4ebp1* (4E-BP1), which encodes a master regulator of cap-dependent translation (Lin et al., 1994; Sonenberg and Hinnebusch, 2009), is induced upon reduction of Rps6 in embryonic limb buds in a p53-dependent manner, suggesting that *Eif4ebp1* may be a p53 transcriptional target gene (Figure 5C-D). Notably, we did not observe transcriptional induction of the closely related homolog, *Eif4ebp2* (Figure 5D), suggesting that this response was specific to *Eif4ebp1*. To examine 4E-BP1 expression at the protein level, given the lower sensitivity of immunoblot analysis compared to RT-qPCR, we dissected out E10.5 *Rps6^lox/+^* and

**Figure 5.**
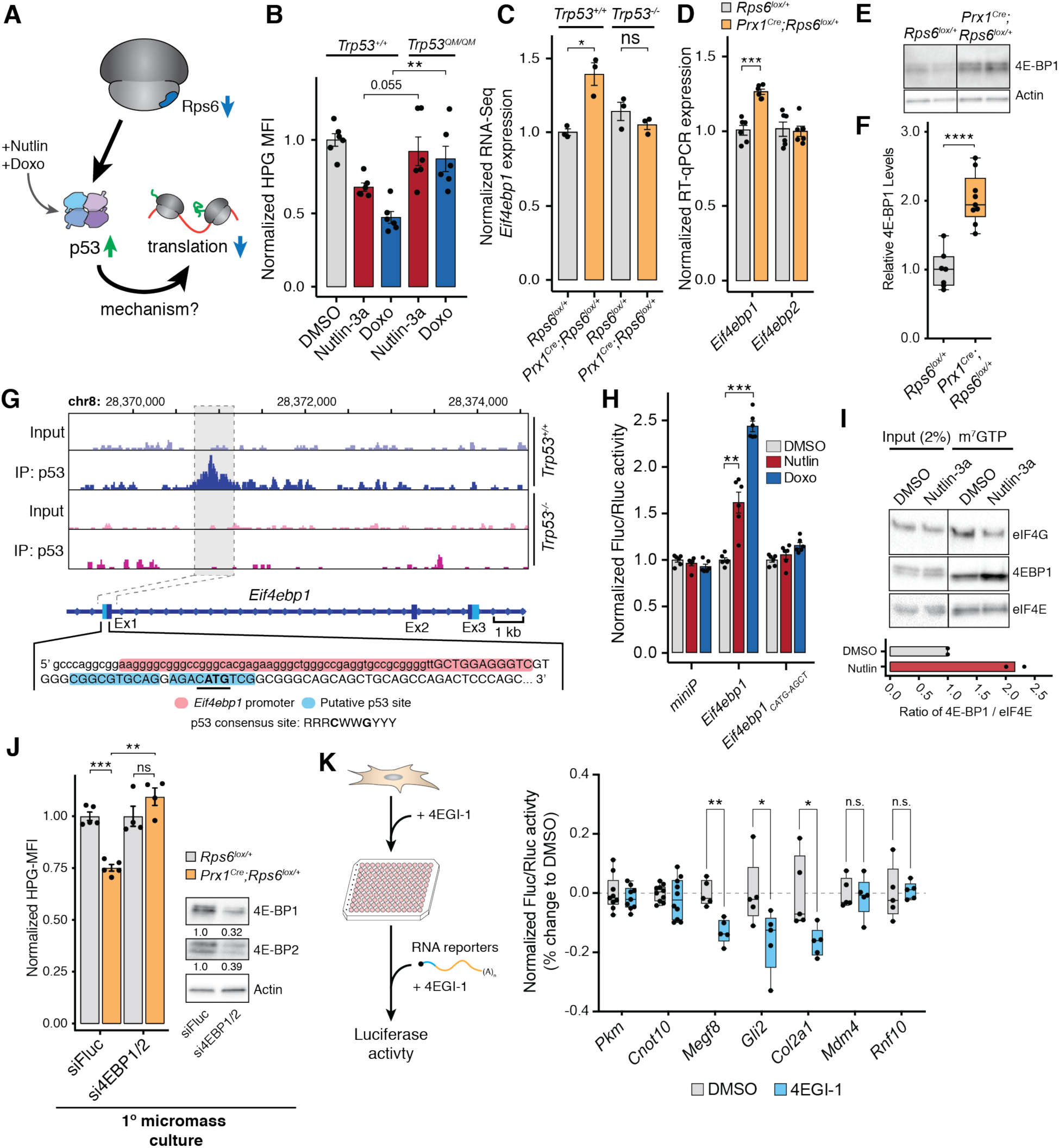
p53 controls translation via transcriptional upregulation of *Eif4ebp1*. (**A**) Potential model of p53-dependent translational control upon *Rps6* haploinsufficiency. (**B**) HPG MFI of MEFs expressing wildtype or transactivation dead p53 (*Trp53^QM^*) treated with Nutlin-3a or doxorubicin normalized to mean of DMSO control expressing wildtype p53. 8 h treatment, *n* = 6. (**C**) *Eif4ebp1* expression from RNA-Seq of E10.5 forelimbs normalized to wildtype (*Rps6^lox/+^;Trp53^+/+^*), *n =* 3. (**D**) RT-qPCR of *Eif4ebp1* and *Eif4ebp2* from E10.5 forelimbs normalized to the geometric mean of housekeeping genes (*Actb, Tbp, Ubb, Exo5, Pkm2, Nadk2*) and wildtype (*Rps6^lox/+^;Trp53^+/+^*), *n =* 6. (**E**) Representative Western blot of 4E-BP1 from E10.5 forelimb mesenchyme cells after removal of the ectoderm layer in *Rps6^lox/+^* and *Prx1^Cre^;Rps6^lox/+^* backgrounds. Complete blot is shown in Figure S13B. (**F**) Relative 4E-BP1 protein levels from *Rps6^lox/+^* and *Prx1^Cre^;Rps6^lox/+^* E10.5 forelimb mesenchyme cells with values normalized to Actin and *Rps6^lox/+^.* Quantification from Figure S13B. *n* = 7 for *Rps6^lox/+^*; *n* = 9 for *Prx1^Cre^;Rps6^lox/+^* (**G**) p53 ChIP-Seq gene track of *Eif4ebp1* locus (Kenzelmann Broz et al., 2013). Transcription start site denoted by capital letters with the *Eif4ebp1* promoter highlighted (red). Putative p53-binding region is highlighted (blue) with the *Eif4ebp1* start codon bolded and p53 core binding sequence is underlined. (**H**) Quantification of Firefly luciferase activity normalized to Renilla luciferase transfection control and to DMSO control of each construct. Plasmid containing either a minimal promoter, the *Eif4ebp1* region, or mutated *Eif4ebp1* region (see Figure S13F) were transfected into NIH3T3 fibroblast cells. Cells were treated with DMSO, Nutlin-3a, or doxorubicin for 8 h; *n =* 6. (**I**) Representative cap-binding assay of NIH3T3 fibroblast cells treated with Nutlin-3a for 8 h. Bottom: ratio of 4E-BP1 to eIF4E in cap-binding assays; *n = 2*. (**J**) **Left**: HPG MFI upon 4E-BP1/2 siRNA treatment in primary *Rps6^lox/+^* and *Prx1^Cre^;Rps6^lox/+^* limb micromass cultures from E10.5 forelimbs. Values normalized to wildtype; *n* = 4. **Right**: Western blot of 4E-BP1 and 4E-BP2 levels in primary *Rps6^lox/+^* limb micromass cultures after siRNA treatment for 16 h. Numbers indicate quantification of proteins normalized to siFluc. (**K**) **Left**: C3H/10T1/2 mesenchymal cells were treated with 4EGI-1 (50 µM) or DMSO for 4 h prior to transfection with luciferase reporter RNAs. **Right**: Firefly/Renilla luciferase activity of RNA reporters transfected into C3H/10T1/2 cells after treatment with 4EGI-1. Activity was normalized to the geometric mean of *Pkm* and *Cnot10* 5’UTR and DMSO. For box-plots, center line, median; box limits, first and third quartiles; whiskers, 1.5x interquartile range; points, outliers. * *P* < 0.05, ** *P* < 0.01, *** *P* < 0.001, **** *P* < 0.0001, two-tailed t-test, unequal variance. For all bar plots, error bars = SEM, * *P* < 0.05, ** *P* < 0.01, *** *P* < 0.001, **** *P* < 0.0001, two-tailed t-test, unequal variance.

*Prx1^Cre^;Rps6^lox/+^* forelimbs and removed the outer unrecombined ectoderm layer before analysis. Accordingly, and in agreement with our RT-qPCR and RNA-seq data, we also observed an increase in 4E-BP1 at the level of protein upon *Rps6* haploinsufficiency (Figure 5E-F; Figure S13B). Moreover, since we did not observe any limb phenotypes or a reduction in protein synthesis upon *Rps6* haploinsufficiency in the AER (Figure 1H-I,N), we asked if *Eif4ebp1* is also induced in this context. Intriguingly, *Eif4ebp1* upregulation was not detected upon Rps6 reduction in the AER (Figure S13C) despite p53 activation in these cells (Figure S3A-B), suggesting that there may be additional tissue-specific regulatory mechanisms governing *Eif4ebp1* expression.

To mechanistically determine whether *Eif4ebp1* is a direct p53 transcriptional target gene, we re-analyzed our previously generated genome-wide p53 ChIP-Seq data (Kenzelmann Broz et al., 2013). This data revealed a ∼0.4 kb region in the first exon of *Eif4ebp1* containing a putative p53 binding site, which was validated by ChIP-qPCR (Figure 5G; Figure S13D). This finding was specific to *Eif4ebp1*, and no candidate p53 binding site was found in the *Eif4ebp2* locus (Figure S13E). When cloned into a luciferase reporter construct (Figure S13F), the ∼0.4 kb *Eif4ebp1* fragment containing a putative p53 binding site was sufficient to drive expression of luciferase upon p53 activation (Figure 5H). Mutation of the core p53 binding sequence (*Eif4ebp1CATG-AGCT*) abolished p53-dependent luciferase activity (Figure 5H), revealing a genuine *cis*-regulatory element responsible for p53-dependent induction of *Eif4ebp1* expression.

The primary function of 4E-BP1 is to repress the major cap-binding protein, eukaryotic initiation factor 4E (eIF4E) (Sonenberg and Hinnebusch, 2009). eIF4E is rate limiting for the translation of select mRNAs, including those with specific sequences or structures within their 5’UTRs. For example, highly structured 5’UTRs are more sensitive to eIF4E dosage as it enhances the unwinding of RNA secondary structures during ribosome scanning by eIF4A (Pelletier et al., 2015). In addition, distinct sequences in the 5’UTR of selective mRNAs such as the cytosine rich motif (CERT) also confer specificity to eIF4E activity (Truitt et al., 2015). Thereby, increased expression of 4E-BP1 inhibits the formation of the eIF4E translation initiation complex and alters the translation of eIF4E-sensitive transcripts. To functionally test whether p53 activation leads to 4E-BP1-mediated cap-dependent translational repression, we first performed cap-binding assays to determine if there is increased binding of 4E-BP1 to eIF4E upon p53-activation. Indeed, we observed an increased ratio of 4E-BP1/eIF4E binding in Nutlin-3a treated cells (Figure 5I). This was accompanied by a concurrent decrease in eIF4G binding, an initiation factor that promotes cap-dependent translation when bound to eIF4E (Figure 5I). Next, we measured whether p53-dependent repression of global protein synthesis requires *Eif4ebp1* expression by depleting 4E-BP in *Rps6^lox/+^* and *Prx1^Cre^;Rps6^lox/+^* primary micromass cultures and performing HPG labeling (Figure 5J; Figure S14A). As 4E-BP1 and 4E-BP2 are functionally redundant, we first knocked-down both proteins to prevent compensation of 4E-BP1 by 4E-BP2 (Bi et al., 2017; Ding et al., 2018). Importantly, 4E-BP1/2 knockdown rescues to a large extent the global protein synthesis repression observed upon Rps6 reduction in *Prx1^Cre^;Rps6^lox/+^* primary micromass cultures (Figure 5J) and in Nutlin-3a treated cells (Figure S14B). This effect was mirrored when 4E-BP1 was knocked-down alone in Nutlin-3a treated cells (Figure S14C).

Our previous results demonstrated that key transcripts involved in limb development are translationally down-regulated in accordance with their 5’UTR features (Figure 4G). Since the 5’UTRs of mRNAs govern their sensitivity to 4E-BP1-eIF4E regulation, we next asked if the 5’UTRs of translationally down-regulated transcripts in our ribosome profiling data were also more sensitive to 4E-BP1-mediated translational regulation. To this end, we took advantage of the small-molecule inhibitor of protein synthesis, 4EGI-1 (eIF4e/eIF4G interaction inhibitor 1), to chemically mimic the effects of 4E-BP1. Mechanistically, 4EGI-1 binds with eIF4E to specifically disrupt association of eIF4G, a large scaffold protein that recruits the 40S ribosomal subunit, while promoting and stabilizing the binding of 4E-BP1 (Sekiyama et al., 2015). We treated C3H/10T1/2 mesenchymal cells with 4EGI-1 prior to transfection with RNA reporter constructs harboring 5’UTRs that were previously demonstrated to be sensitive to RP haploinsufficiency in either a p53-dependent or -independent manner (Figure 4D,G; Figure 5K). We found that reporter constructs containing 5’UTRs whose translational regulation was p53-dependent (*Megf8, Gli2, Col2a1*) to be more sensitive to 4EGI-1 treatment compared to p53-independent 5’UTRs (*Mdm4, Rnf10*) or control 5’UTRs that do not change in TE in our ribosome profiling experiment (*Pkm, Cnot10*) (Figure 5K). These data further show that 4E-BP1-mediated translation regulation contributes to the selective p53-dependent translational changes observed upon *Rps6* haploinsufficiency.

To further understand if 4E-BP1 upregulation is a general phenomenon associated with depletion of core RPs, we knocked down RPs commonly mutated in ribosomopathies such as DBA and 5q-myelodysplastic syndrome, *Rps19* and *Rps14*, in primary limb micromass cultures and assessed p53 activation and *Eif4ebp1* expression. As with *Rps6* haploinsufficiency, we found that p53 is activated upon depletion of *Rps19* or *Rps14*, which coincides with an increase in *Eif4ebp1* expression (Figure 6). Together, these results suggest a common pathway via the p53-4EBP1-eIF4E axis by which selective changes in translation occur upon ribosome perturbation, including in the context of RPs directly mutated in ribosomopathies.

**Figure 6.**
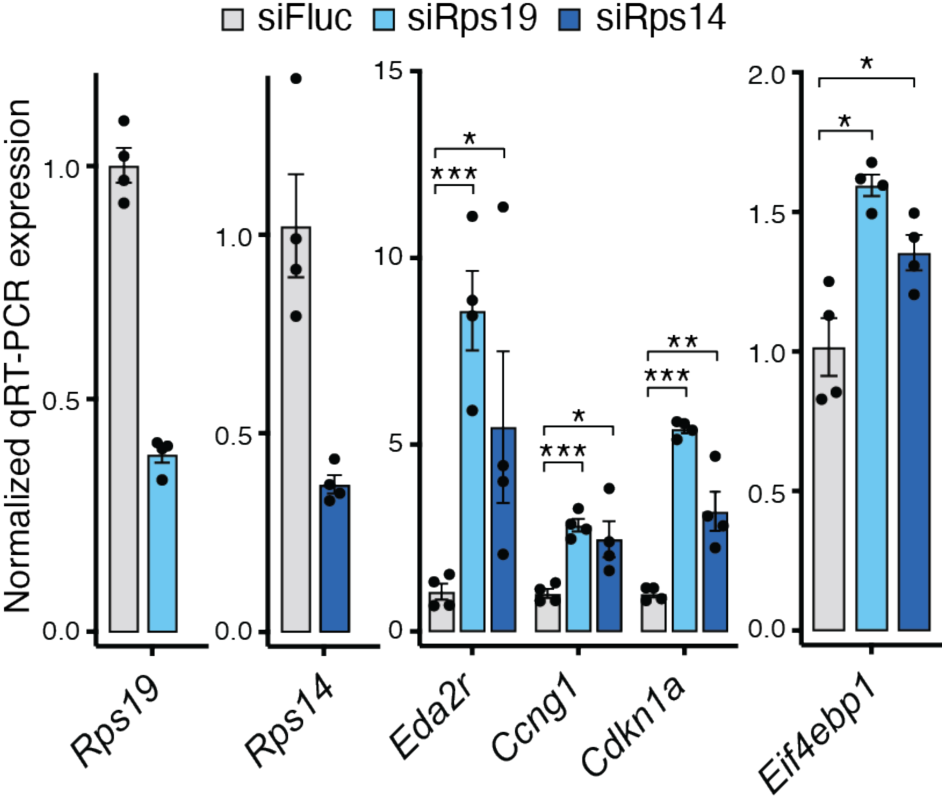
Depletion of ribosomal proteins commonly mutated in ribosomopathies induces *Eif4ebp1* expression. Relative mRNA expression of p53 target genes and *Eif4ebp1* in primary limb micromass cultures after siRNA knockdown of *Rps19* and *Rps14*. *Rps19* and *Rps14* levels upon knockdown are also shown. Expression was normalized to the geometric mean of *TBP* and *NupL1* and then to siRNA control. *n = 4*. Error bars = SEM, * *P* < 0.05, ** *P* < 0.01, *** *P* < 0.001, **** *P* < 0.0001, two-tailed t-test, unequal variance.

## DISCUSSION

Ribosomopathies caused by haploinsufficiency of essential ribosomal proteins present with a variety of phenotypes including craniofacial, digit, and limb abnormalities, as well as selective impairment of erythroid differentiation. How such phenotypes, and in particular congenital birth defects, are produced from mutations in essential ribosome components has been an outstanding mystery for the last two decades. Studies investigating the role of p53 in ribosomopathies have found that p53 activation contributes to phenotypes in many model systems, including in zebrafish, mouse, and human cells (Danilova et al., 2008; Dutt et al., 2011; Jones et al., 2008; McGowan et al., 2008, 2011; Sulic et al., 2005). However, how p53 activation can shape complex, tissue selective phenotypes such as radial aplasia and other congenital birth defects is poorly understood. The few studies that have suggested that translation dysregulation plays a role in ribosomopathies typically hypothesized that these translational changes occur directly from impaired ribosome function, and do not reconcile these findings with the known role of p53 in producing phenotype (Horos et al., 2012; Khajuria et al., 2018; Ludwig et al., 2014). Such studies have relied on *ex vivo* models and used strong knockdowns of RPs that likely reduce RP levels much lower than what would be physiologically expected in an RP haploinsufficient disease. It has therefore been difficult to capture a physiologically relevant change in translation control underlying *in vivo* tissue phenotypes. Thus, while studies of p53-dependent or - independent mechanisms have typically been considered to be mutually exclusive processes that underlie ribosomopathies, our work surprisingly demonstrates that they are interconnected in an *in vivo* model of RP haploinsufficiency. In particular, we demonstrate an unexpected role of p53 in mediating much of the translation program downstream of RP haploinsufficiency. We show that p53 transcriptionally induces the expression of the translation inhibitor 4E-BP1, which is the main negative regulator of eIF4E-mediated cap-dependent translation (Lin et al., 1994; Sonenberg and Hinnebusch, 2009), through a p53-binding element in the first exon of the *Eif4ebp1* gene locus. Thereby, translational specificity to gene expression upon RP haploinsufficiency can arise from an intermediary pathway, the p53-4E-BP1-eIF4E axis, which becomes activated and links RP haploinsufficiency to selective changes in cap-dependent translation.

Our discovery that p53-mediated regulation of 4E-BP1 underlies many of the translational changes upon RP haploinsufficiency also suggests mechanisms by which tissue specificity can emerge in ribosomopathies. Tissue specific phenotypes could emerge either through 4E-BP1-mediated selective translation of key developmental transcripts within tissues or through differential expression and activity of 4E-BP1 itself among different tissues. The 4E-BP1-eIF4E axis is a rate-limiting component of translation initiation (Duncan et al., 1987). In particular, certain transcripts with specific sequences and structures in their 5’UTRs are especially sensitive to eIF4E mediated translational regulation (Pelletier et al., 2015; Truitt et al., 2015). As a result, p53-mediated induction of 4E-BP1 may lead to a translational program in which perturbed translation of select transcripts, which are differentially sensitive to eIF4E activity, contribute to tissue specific phenotypes. For example, in the limb upon *Rps6* haploinsufficiency, we observe p53-dependent translational repression of key limb developmental transcripts that correlate with the observed limb development phenotype. In addition, we also observed that differential 4E-BP1 expression and response among different tissues may also contribute to cell and tissue specific phenotypes upon RP haploinsufficiency. For example, we demonstrate that although conditional *Rps6* haploinsufficiency led to p53 activation in both the limb mesenchyme and AER, reduction of Rps6 led to 4E-BP1 upregulation and subsequent translation repression associated with a selective phenotype only in the limb mesenchyme. As a result, future studies to characterize tissue specific 4E-BP1 expression and to investigate the mechanisms underlying such selectivity would provide further insight into tissue specific phenotypes in ribosomopathies.

Our findings demonstrating a role for p53-mediated induction of 4E-BP1 in translational control downstream of RP haploinsufficiency also suggest possible therapeutic strategies to treat ribosomopathies. Treatment with *l*-leucine, which upregulates translation via activation of the mTORC1 pathway, has been shown to rescue developmental phenotypes in some models of ribosomopathies (Jaako et al., 2012; Payne et al., 2012). Given that our studies demonstrate that *in vivo* phenotypes upon RP haploinsufficiency are driven at least in part by translational perturbation via the p53-4E-BP1-eIF4E axis, more potent activators of cap-dependent translation and eIF4E activity may serve as enticing candidates to treat ribosomopathies. Furthermore, despite activation of the tumor suppressor p53 observed in ribosomopathies such as DBA (Dutt et al., 2011), patients with DBA also surprisingly present with a higher incidence of cancer including leukemia and osteogenic sarcoma (Vlachos et al., 2012). In the context of our findings showing that p53 is a master regulator of protein synthesis, one explanation for how this paradoxical increase in cancer risk could occur would be that these cells lose their ability to repress eIF4E-mediated translational control via 4E-BP1 upon inactivation of p53. Indeed, increased eIF4E activity has been shown to promote cellular transformation and tumor formation (Furic et al., 2010; Mamane et al., 2004; Ruggero et al., 2004). In fact, inactivating mutations in p53 occur frequently in cancer (Kastenhuber and Lowe, 2017), and in these cancers, 4E-BP1 expression and protein synthesis may be dysregulated. Therefore, inhibitors of cap-dependent translation may also serve as enticing new candidates to treat cancers characterized by loss of p53.

Although most of the translational changes observed in *Prx1^Cre^;Rps6^lox/+^* mice were dependent on p53, we also observed a small number of p53-independent translational changes in our ribosome profiling data that were validated by polysome fractionation. These changes included translational upregulation of nearly all ribosomal proteins and translational repression of the p53 negative regulator *Mdm4*. We hypothesize that these changes represent possible post-transcriptional quality control mechanisms that allow cells to quickly respond to ribosome dysfunction. For example, translational upregulation of RPs upon haploinsufficiency of a specific RP would allow for increased RP production that may help preserve ribosome homeostasis and mitigate the effects of RP depletion. Furthermore, translational repression of *Mdm4*, a negative regulator of p53, may serve as a quick post-transcriptional quality control sensor to activate p53 in the context of RP haploinsufficiency.

Ribosome biogenesis is one of the most energetically expensive programs in the cell. By coupling alterations in the expression of essential RPs directly to cap-dependent translation via 4E-BP1, p53 may ensure that a modified cellular translatome continues to support cellular growth and survival under suboptimal conditions. This is likely highly dependent on the nature of the transcripts expressed within specific cellular and/or tissue contexts and would result in translational downregulation of specific mRNAs that manifests in tissue specific phenotypes. Interestingly, recent findings suggest that the *Drosophila* Minute phenotype, which is caused by RP haploinsufficiency and is characterized by short bristles and a reduction in global protein synthesis, also largely occurs indirectly as a result of a regulatory response to RP haploinsufficiency (Kongsuwan et al., 1985; Lee et al., 2018). The phenotype and translational changes were driven by the bZip-domain protein Xrp1 rather than by a direct decrease in ribosome levels (Lee et al., 2018). As opposed to Xrp1 in *Drosophila*, we show that p53 itself drives the majority of the translational changes in an *in vivo* mammalian model of RP haploinsufficiency. Therefore, a direct link between p53 and 4E-BP1-eIF4E activity established in this study suggests that characterization of p53-dependent remodeling of the translational landscape is likely to yield greater insights and novel therapeutic strategies to target ribosomopathies.

## AUTHOR CONTRIBUTIONS

M.B. and G.C.T. conceived, and M.B. supervised the project; G.C.T., C.H.K., and M.B. designed the experiments; G.C.T., C.H.K., P.S.K., H.D.R., M.E.B., L.S., and O.Z. performed mouse embryo experiments and phenotyping; C.H.K. performed cell culture experiments; G.C.T., C.H.K., C.M.F., and

N.R. performed metabolic labeling experiments; G.C.T. performed and analyzed ribosome profiling experiments; G.C.T. and C.H.K. performed RT-qPCR experiments; C.H.K. performed Western blotting experiments and reporter assays; N.R., M.E.B., and L.D.A. guided experiments related to p53; C.M.F. and D.R. guided experiments related to OP-Puro labeling and characterization of global protein synthesis changes; G.C.T. and C.H.K. performed all other experiments; M.B., G.C.T., and C.H.K. wrote the manuscript with input from all the authors.

## ACKNOWLEDGMENTS

The authors would like to thank Kotaro Fujii (Stanford), Katrina Cabaltera, Jing Huang (Stanford), Naomi Genuth (Stanford), and David Mahoney (Stanford) for their experimental contributions as well as the Stanford FACS facility. The authors would also like to thank George Thomas (IDIBELL) for generously providing the *Rps6^lox^* mice, and members of the Barna lab for critical feedback and discussion. **Funding:** Supported by New York Stem Cell Foundation grant NYSCF-R-I36 (M.B.), New York Stem Cell Robertson Investigator (M.B.), NIH grant 1R01HD086634 (M.B.), Alfred P. Sloan Research Fellowship (M.B.), Pew Scholars Award (M.B.), NIH grant R35CA242986 (D.R.), March of Dimes Foundation grant no. 6-FY15-189 (L.D.A.), NIH R35 grant CA197591 (L.D.A.), Paul and Daisy Soros Fellowships for New Americans (G.C.T), Stanford Medical Scientist Training Program (G.C.T.), Canadian Institutes for Health Research Fellowship (C.H.K. and O.Z.), Stanford Dean’s Fellowship (C.H.K.), Helen Hay Whitney Foundation Fellowship (O.Z.), NIH/NIDDK grant K08DK119561 (C.M.F.), National Institutes of Health Developmental Biology Training Grant 5T32GM007790-41 (H.D.R.), Campini Foundation and the University of California, San Francisco, Department of Pediatrics K12 (Award 5K12HDO72222-05) (C.M.F.), and the Jane Coffin Childs Fund Postdoctoral Fellowship (M.E.B.). **Data and materials availability:** All data are available in the manuscript or the supplementary materials. Data that support the findings of this study have been deposited in the Gene Expression Omnibus under accession number GSE135722.

## MATERIALS AND METHODS

### Mouse husbandry

All animal work was reviewed and approved by the Stanford Administrative Panel on Laboratory Animal Care (APLAC). The Stanford APLAC is accredited by the American Association for the Accreditation of Laboratory Animal Care (AAALAC). All mice used in the study were housed at Stanford University. Mice were maintained on mixed backgrounds and the following previously described alleles were used: *Rps6^lox^* (Volarevic et al., 2000), *Prx1^Cre^* (Logan et al., 2002), *Msx2^Cre^* (Sun et al., 2000), *Tsc2^lox^* (Hernandez et al., 2007), *Trp53^null^* (Jacks et al., 1994), and *Rosa^mTmG/+^* (Muzumdar et al., 2007). *Rps6^lox^* mice were generously provided by George Thomas (IDIBELL). *Prx1^Cre^*, *Msx2^Cre^*, and *Tsc2^lox^* mice were obtained from the Jackson Laboratories. Genotyping was performed using standard PCR protocols using primers described in aforementioned publications or on the Jackson Laboratories website. Common *Cre* primers were used to genotype *Cre* lines (http://mgc.wustl.edu/protocols/pcr_genotyping_primer_pairs). The following primers were used for *Rps6^lox^* genotyping: Rps6lox_1 GCTTCTACTTCTAAGTCTGAGTCCAGTC, Rps6lox_2 TCCTGCCGAGAAAGTATCCATCATG, and Rps6lox_3 CTGCAGCCTTTTCTTTTAGCATACCTG. Mice for a given experiment were colony-matched. Mouse embryos for a given experiment were somite matched. Mouse embryos were harvested at E10.5, E11.5, E12.5, E13.5, and E17.5.

### Cell Lines

Primary MEFs were derived from E13.5 mouse embryos using standard protocols and cultured under standard conditions in DMEM supplemented with 2 mM L-glutamine and 10% fetal bovine serum (FBS). Cells were used for experiments at passages 2-5. NIH3T3 and C3H/10T1/2 cells were purchased from the American Type Culture Collection and grown under standard conditions in DMEM supplemented with 10% FBS and 2 mM L-glutamine. Cells were passaged 1:6 roughly every 2-3 days. All cell lines used in this study were mycoplasma-free.

### Skeleton and cartilage staining

Bone and cartilage staining of E17.5 embryos were performed as per standard methods. Skeletal images were acquired using a Zeiss SteREO Discovery.V8 microscope and a Zeiss AxioCam MRc5 camera with an external standard as scale. Skeletal elements were measured using the ImageJ Fiji plug-in with genotypes blinded during measurement.

### Separation of limb ectoderm and mesenchyme

Forelimb buds of E10.5 or E11.5 embryos were dissected in 1×PBS. Dissected limb buds were then digested in 3% trypsin in 1×PBS for 25 min at 4°C with gentle shaking. Trypsin digestion was quenched by moving the limb buds to 10% FBS in PBS. The limb buds were then gently vortex for 15 seconds to dislodge the mesenchymal and ectodermal layers. The ectodermal jackets were further removed from the individual limb buds under a dissecting microscope. Samples were stored at -80°C until use.

For OP-Puro labeling, dissected ectoderm was dissociated from the mesenchyme layer of E11.5 forelimb buds by incubating in 1% trypsin at room temperature for 20 min. The ectoderm was then separated from the mesenchyme layer and resuspended in DMEM/F12, no phenol red, 10% FBS and labeled with OP-Puro (20 µM) as described below.

### *In situ* hybridization

*In situ* hybridization for *Sox9* at E12.5 was performed as per standard protocols (Lufkin, 2007) with modifications to achieve good probe penetration and signal. Digoxigenin (DIG) labeled RNA probes were generated from *XhoI* linearized T3 promoter *Sox9* plasmid (gift from Chi-Chung Hui, University of Toronto). Fixed and rehydrated embryos were treated with 20 µg/mL proteinase K in PBS+0.1% Tween-20 (PBST) for 35 minutes at room temperature. After post-fixation and subsequent washes, embryos were incubated overnight at 70°C with hybridization mix containing 0.5 µg/mL denatured DIG-labeled probe. After washes and pre-blocking, embryos were incubated overnight at 4°C with 1:2000 alkaline phosphatase-conjugated anti-DIG antibodies (Sigma/Roche, 11093274910). After washes and prior to development, embryos were washed 2x20 min with 100 mM NaCl, 100 mM Tris-HCl pH 9.5, 50 mM MgCl2, 0.1% Tween-20, 2 mM levamisole. Embryos were then developed with BM Purple AP substrate (Sigma/Roche, 11442074001) at 4°C for >1 day until adequate signal to noise ratio was achieved. The reaction was then subsequently inactivated, post-fixed, and stored. Embryos were then imaged with a Leica MZ16 FA microscope with a Leica DFC480 camera with an external standard used for scale.

### Vectors and *in vitro* transcription

For generation of 5’UTR reporter constructs, sequences were obtained from ENSEMBL and isoforms were chosen based on alignment to the RNA sequencing results presented herein. 5’UTR gene fragments were synthesized (Twist Biosciences) and cloned immediately upstream of a Firefly luciferase gene encoded in pGL3-FLB (Leppek *et al*. unpublished). Each 5’UTR-*Fluc* construct was PCR amplified using primers flanking the 5’UTR and *Fluc* gene and incorporating a T7 RNA polymerase promoter at the 5’ end. PCR amplicons were column purified (NEB, T1030) and used as a template for *in vitro* transcription reactions with a mMESSAGE mMACHINE® T7 Transcription Kit (Ambion, AM1344) followed by the addition of a poly(A) tail using a Poly(A) Polymerase Tailing Kit (Lucigen, PAP5104H). *In vitro* synthesized capped and poly(A) tailed RNA was purified using PureLink RNA Mini Columns (Thermo Fisher, 12183020). RNA integrity was confirmed via denaturing agarose gel electrophoresis.

For testing p53 transcriptional activity, a 423 bp gene fragment of *Eif4ebp1* was amplified from genomic DNA of primary mouse embryonic fibroblasts and cloned in lieu of the minimal promoter sequence within pGL4.23[*luc2/miniP*]. All constructs were sequence verified. **siRNA-mediated knockdowns.** NIH3T3 cells were plated at a density of 1.0×10^5^ cells/well in a 6-well plate and allowed to attach for at least 6 h at 37°C. Cells were washed with 1×PBS before being transfected with 25 nM of non-targeting control siRNA or siRNA target against 4E-BP1 and 4E-BP2 (Dharmacon) using Dharmafect Reagent I (Dharmacon, T-2001) as per manufacturer’s protocol. Cells were incubated for 16 h at 37°C in antibiotic-free media. Following incubation, the siRNA-containing media was replaced with complete DMEM media (as described above) and cells were treated with either DMSO, Nutlin-3a, or Doxorubicin.

For siRNA treatment in micromass limb cultures, limbs from E11.5 embryos were dissected and cells dissociated as described. Subsequently, 1.5×10^6^ cells were reverse transfected with 25 nM siRNA as per manufacturer’s protocol using Dharmafect I (Dharmacon, T-2001) in Limb micromass media and plated in a 96-well plate. Cells were incubated for 20 h before use in subsequent experiments. Note that siRNA knockdowns of *Rps19* and *Rps14* were performed in micromass cultures derived from wildtype *C57BL/6J* mice.

### RT-qPCR quantification of RNA expression

RNA from embryonic limbs or cell culture was extracted with TRIzol (Thermo Fisher, 15596026) as per manufacturer’s protocol followed by DNase I treatment for 30 min at 37°C. RNA was then purified via Zymo RNA Clean & Concentrator columns (Zymo Research). Subsequently, 100 ng of RNA was used for first strand cDNA synthesis using the iScript Supermix (Bio-rad, 1708841). cDNA was diluted 18-fold and 4 µL was used in a reaction for SYBR green detection with SsoAdvanced SYBR Green supermix (Bio-Rad, catalog no. 1725270) on a CFX384 machine (Bio-rad). All primer sequences are available in Table S2.

### Western blots

Equal amounts of protein were resolved on either a 15% or a 4-20% Tris-glycine gradient SDS-PAGE gel and transferred to a polyvinylidene difluoride Immobilon-FL membrane (PVDF; Bio-rad). Membranes were blocked for 30 min at room temperature with 5% BSA in TBST (50 mM Tris, 150 mM NaCl, 1% Tween-20, pH 7.4). Blots were incubated for 24 h at 4°C with the following primary antibodies: mouse anti-GAPDH (Cell Signalling, 97166), mouse anti-actin (Cell Signalling, 3700S), rabbit anti-Rps6 (Cell Signalling, 2217L), rabbit anti-4E-BP1 (Cell Signalling, 9644S), rabbit anti-phospho-p70 S6 Kinase (Cell Signalling, 9205S), rabbit anti-p70 S6 Kinase (Cell Signalling, 9202S), rabbit anti-phopsho-ULK1(Ser757) (Cell Signalling, 6888T), rabbit anti-ULK1 (Cell Signalling, 8054T), rabbit anti-p53 (Leica Biosystems, CM5), rabbit anti-TSC2 (Cell Signalling, 4308T), rabbit anti-eIF4G (Cell Signalling, 2498S), mouse anti-eIF4E (BD Transduction, 610269). All antibodies were used at a dilution of 1:1000 in 5% BSA-TBST, unless stated otherwise. Membranes were washed 3 times for 5 min in TBST before incubation for 1 h at room temperature with secondary antibodies: donkey anti-mouse (1:5000; GE Healthcare, NA931-1ML), or donkey anti-rabbit (1:5000; GE Healthcare, NA934-1ML) coupled to Horseradish Peroxidase. Membranes were then washed 3 times for 5 min in TBST before detection using Clarity Western ECL Substrate (Bio-rad, 170-5061) and imaging on a ChemiDoc MP (Bio-rad, 17001402). All blots were quantified using ImageJ v2.0.0.

### Micromass cultures and RNA transfection luciferase assays

E11.5 mouse limbs were harvested in DMEM/F12, no phenol red, 10% FBS, pen/strep. Cells were then dissociated in Dissociation Buffer (1% trypsin in HBSS without Ca^+2^ or Mg^+2^) and incubated for 30 min at 37°C. Single-cell suspensions were obtained by quenching the dissociation with Limb Media (DMEM/F12 HEPES, 10% FBS) and passing through a cell strainer. Cells were then pelleted and resuspended in Limb Media. Next, 1.0×10^5^ cells were co-transfected with capped and poly(A)-tailed Fluc (100 ng) and Rluc reporter RNAs (10 ng) using Lipofectamine 2000 (Thermo Fisher, 11668-019) before seeding into a single well of a 96-well Nunclon Delta Microplate (Thermo Fisher, 167008). Cells were incubated for 6 h at 37°C before being harvested in 1× Passive Lysis Buffer and subjected to a Dual-Luciferase Reporter Assay System as per manufacturer’s instructions (Promega, E1980) on a GloMax-Multi plate reader (Promega). Resulting luciferase activities were first normalized to the geometric mean of *Pkm* and *Cnot10* 5’UTR activities and then to wildtype micromass cultures.

### DNA transfection luciferase assays

NIH3T3 cells were seeded at a density of 1.0×10^5^ cells/well in a 6-well plate and incubated at 37°C overnight. The following day cells were co-transfected with 2 µg of Fluc plasmid DNA and 20 ng of Rluc control plasmid (pRL) per well using Lipofectamine 2000 (Thermo Fisher, 116688-019) and incubated for 16 h at 37°C. Cells were then treated with either DMSO, Nutlin-3a (10 µM), or Doxorubicin (0.2 µg/mL) for 8 h. Cells were harvested and lysed in 50 µL of 1×Passive Lysis Buffer. Lysate was cleared of debris by centrifugation at 10,000 RCF for 5 min, and 20 µL of resulting supernatant was used to measure luciferase activity with the Dual-Luciferase Reporter Assay System (Promega, E1980) on a GloMax-Multi plate reader (Promega).

### 4EGI-1 treatment of 10T1/2 cells and luciferase assays

10T1/2 cells were seeded at a density of 4.0×10^5^ cells/well in a 96-well plate and incubated at 37°C overnight. The next day, cells were treated with 50 µM 4EGI-1 (Sigma, 324517) or DMSO in DMEM supplemented with 10% FBS and 2 mM L-glutamine and incubated for 4 h at 37°C. Following treatment, cells were co-transfected with capped and poly(A)-tailed Fluc (100 ng) and Rluc reporter RNAs (10 ng) using Lipofectamine 2000 (Thermo Fisher, 11668-019). Cells were then re-treated with 50 µM 4EGI-1 or DMSO and incubated for a further 4 h at 37°C. After a total of 8 h of treatment (4 h prior to transfection and 4 h after transfection), cells were harvested in 1× Passive Lysis Buffer and luciferase activities were measured using a Dual-Luciferase Reporter Assay System as per manufacturer’s instructions (Promega, E1980) on a GloMax-Multi plate reader (Promega). Resulting luciferase activities were normalized to the geometric mean of *Pkm* and *Cnot10* 5’UTR activities and then to DMSO treated cells.

### P53 transactivation domain mutated MEFs

MEFs were derived from previously generated knock-in mice expressing p53 wild-type (*Trp53^LSL-WT/LSL-WT^*) or *Trp53^QM/QM^* (*Trp53^LSL-25,26,53,54/LSL-25,26,53,54^*) mutant(Brady et al., 2011). In these MEFs, expression of both *Trp53* alleles is silenced through upstream transcriptional stop element flanked by *loxP* recombination sites. To reactivate p53 expression, MEFs were infected with adenoviruses expressing Cre recombinase (Ad5 CMV-Cre) or empty virus (Ad5-CMV-Empty) as control at an approximate multiplicity of infection of 100. Viruses were purchased from University of Iowa Viral Vector Core Facility. Infected cells were cultured for 48 hours at 5% CO2, 37°C before initiating drug treatments and HPG assay. Cells were treated with 0.2 μg/ml doxorubicin or 10 µM Nutlin-3a for indicated times (see below). High efficiency of recombination (>90% cells) was confirmed by immunostaining of p53. P53 stabilization upon doxorubicin and Nutlin-3a treatment was confirmed by Western blotting using p53 CM5 antibody (Leica Biosystems).

### Chromatin immunoprecipitation qPCR

MEFs were grown in DMEM containing 10% FCS and seeded at 7 x 10^6^ cells per 10 cm dish one day prior to the ChIP experiment. After treatment with 0.2 μg/ml doxorubicin for 6h, cells were harvested to prepare chromatin for immunoprecipitation using p53 polyclonal antibodies (NCL-L-p53-CM5p; Leica Biosystems). ChIPs were performed essentially as described previously (Kenzelmann Broz et al., 2013). Chromatin-immunoprecipitated DNA was quantified by qPCR using SYBR Green and a 7900HT Fast Real-Time PCR machine (Applied Biosystems) and primers specific for *Eif4ebp1*. The signals obtained from the ChIP were analyzed by the percent input method.

### Cap-binding assays

NIH3T3 cells were seeded at a density of 1.0×10^6^ in 10 cm tissue culture plates and incubated overnight at 37°C. Cells were then treated with DMSO or Nutlin-3a (10 µM) for 8 h. Subsequently, cells were harvested and washed twice with ice cold 1×PBS before being lysed for 30 min on ice with occasional agitation in Cap Lysis Buffer (10 mM Tris-HCl pH 7.5. 140 mM KCl, 4 mM MgCl2, 1 mM DTT, 1 mM EDTA, 1% Nonidet P-40) supplemented with protease and phosphatase inhibitor for 30 min on ice with occasional agitation. Lysates were cleared via centrifugation at 10,000 RCF for 15 min at 4°C. Resulting supernatants were collected and protein concentration was determined using a BCA assay (Thermo Fisher). 250 µg of protein was pre-cleared using 50 µL of agarose beads (Jena Biosciences, AC-100S) in Cap Lysis Buffer for 30 min at 4°C with rocking. Next, the beads were pelleted at 1000 RCF and the supernatant was added to 50 µL of γ-aminophenyl-m7GTP (C10-spacer)-agarose beads (Jena Biosciences, AC-155S) in Cap Lysis Buffer overnight at 4°C with rocking. Supernatant was removed and the beads were washed twice with Cap Lysis Buffer followed by 1×PBS. Beads were then resuspended in 100 µL of 1× SDS loading buffer and boiled for 5 min. 25 µL of each sample was used for SDS-PAGE and Western blot analysis for eIF4E, eIF4G, and 4E-BP1.

### Proliferation and apoptosis assays

Cell proliferation and apoptosis levels were measured using EdU (5-ethynyl-2′-deoxyuridine) incorporation and Annexin V staining, respectively. Briefly, 1.0×10^5^ NIH3T3 cells were seeded in a 6-well plate and incubated overnight at 37°C. The following day cells were treated with DMSO or Nutlin-3a (10 µM) for 4 or 8 h. For proliferation measurements, one hour before incubation was complete, cells were treated with 10 µM EdU using a Click-iT EdU Alexa Fluor 488 flow Cytometry Assay Kit (Thermo Fisher, C10425) and processed as per manufacturer’s instructions. For apoptosis measurements, cells were harvested, washed twice with Cell Staining Buffer, and labeled using a FITC Annexin V and propidium iodide staining kit as per manufacturer’s protocol (BioLegend, 640914). Both EdU- and Annexin V-labeled samples were analyzed on a LSRII flow cytometer (BD Biosciences) using software packages CellQuest and FlowJo v10.

For whole mount cell death assays, dissected embryos were incubated with 1 mL of 5 µM LysoTracker Red DND-99 (Invitrogen) in HBSS for 45 min at 37°C. Embryos were washed 4X with HBSS and then fixed in 4% paraformaldehyde overnight at 4°C. Embryos were washed with PBS + 0.1% Tween-20 and then dehydrated stepwise into 100% methanol in which they were stored at -20°C. Prior to imaging, they were rehydrated stepwise into PBS + 0.1% Tween-20 and imaged on a fluorescence microscope.

### Immunofluorescence on cryosections

Immunofluorescence on cryosectiosns was performed per standard protocols. Mouse embryos were dissected and fixed in ice-cold 4% PFA for 1 hour, washed four times with PBS + 0.1% Tween-20, equilibrated in 30% sucrose 0.1M potassium phosphate buffer, pH 7.4, embedded in O.C.T., and stored at -80°C. Embryos were sectioned transversely into 12 um thick sections using a Leica cryostat, and frozen on slides at -80°C until use. For immunofluorescence staining, all washes and incubations were done using blocking buffer (PBS with 1% goat serum and 0.1% Triton X-100). Sections were blocked for 1 hr in blocking buffer, incubated in primary antibody overnight at 4C, washed three times, incubated for 1 hr at room temperature with secondary antibody, and washed three more times before mounting. Primary antibodies were used at the following concentrations: 1:200 Leica anti-p53 (CM5P-L), 1:500 Sigma anti-phospho-Histone H3 (06-570), 1:600 Cell Signalling Technology Cleaved Caspase-3 (9661S). Goat anti-rabbit AF568 secondary from Life Technologies (A11036) was used for all slides at 1:500. Slides were imaged using a Zeiss EC 10x Plan-Neofluar Ph1 objective (NA = 0.3), and acquired using a CSU-X1 UltraVIEW Spinning disc with a Hamamatsu EM-CCD camera and Volocity software. Slides comparing the same antibodies were imaged using consistent imaging settings and processing. For phospho-Histone H3 quantification, total cell count in the limb bud from multiple embryos was quantified from DAPI staining with Spot Counter in ImageJ, and pHistone3 positive cells were counted manually. Images are representative of average percentages of pHistone3 cells in Prx and WT sections.

### Measurement of global protein synthesis

Briefly, O-propargyl-puromycin (OP-Puro) labeling of embryonic limbs was done as follows. E10.5 embryos were dissected in filming media (DMEM/F12, no phenol red, 10% FBS). Forelimbs were removed and dissociated in Dissociation Buffer (1% trypsin in HBSS without Ca^+2^ or Mg^+2^) for 15 min at 37°C. Resuspended forelimb cells were used for downstream labeling and analysis. For OP-Puro or L-homopropargylglycine (HPG) labeling of cultured cells, primary MEFs or NIH3T3 cells were seeded at a density of 1.0×10^5^ cells/well in a 6-well plate and allowed to attach overnight. The following day, cells were treated with DMSO, Nutlin-3a (10 µM), or Doxorubicin (0.2 µg/mL) before metabolic labeling. For OP-Puro incorporation, cells were labeled with 20 µM of OPP in DMEM plus drug for 30 min at 37°C. For HPG labeling, two hours before each timepoint, cells were methionine starved in Met dropout media (DMEM supplemented with 10% dialyzed FBS, 25 mM L-cysteine, 2 mM L-glutamine, no methionine) for 45 min at 37°C. Cells were then labeled with 50 µM HPG (Thermo Fisher, C10186) or 50 µM L-Met (control) for 1 h at 37°C. Following metabolic labeling, cells were harvested and washed with twice 1×PBS. Cell pellets were resuspended in Zombie Violet Live-Dead Stain (1:500 in PBS; BioLegend, 423113) and incubated for 15 min in the dark. Cells were then washed with Cell Staining Buffer (0.1% NaN3, 2% FBS in HBSS) before being fixed in 1% PFA for 15 min on ice. Subsequently, cells were permeabilized overnight at 4°C in Perm Buffer (0.1% Saponin, 0.1% NaN3, 3% FBS in PBS). The next day, cells were washed twice with Cell Staining Buffer (without 0.1% NaN3), labeled with an Alexa Fluor 555 Picolyl Azide dye (Thermo Fisher, C10642) and incubated for 30 min at room temperature in the dark. Labeled cells were washed and resuspended in Cell Staining Buffer before being analyzed on a LSRII flow cytometer (BD Biosciences) and using software packages CellQuest and FlowJo v10.

### Sucrose density gradients and RT-qPCR

E10.5 mouse embryonic limbs were harvested in cold HBSS (Thermo Fisher, 14025-076) containing 100 μg/ml cycloheximide (CHX). Limb pairs from 3-4 embryos of a given genotype were pooled and lysed via vigorous pipetting and 30 minute incubation at 4°C in 175 μL lysis buffer (20 mM Tris-HCl pH 7.5, 150 mM NaCl, 15 mM MgCl2, 1 mM DTT, 8% glycerol, 1% Triton X-100, 100 μg/ml CHX (Sigma-Aldrich, C7698-1G), 200 U/mL SUPERase RNase Inhibitor (Thermo Fisher, AM2696), 20 U/ml Turbo DNAse (Thermo Fisher, AM2238), and 1X Halt Protease and Phosphatase Inhibitor Cocktail (Thermo Fisher, 78442). Samples were centrifuged at 1,300xg, 5 min and 10,000xg, 5 min to remove cell debris. Samples were then layered onto a 15-45% sucrose gradient or 25-50% sucrose gradient containing 20 mM Tris-HCl pH 7.5, 100 mM NaCl, 15 mM MgCl2, and 100 μg/ml CHX. 15-45% gradient was made on a Biocomp Model 108 Gradient Master. 25-50% gradient was made by sequentially freezing at -80°C a 5-step gradient (50%, 43,75%, 37.5%, 31.25%, 25% sucrose). Samples layered on gradients were spun on a Beckman SW-60 rotor at 35,000 RPM, 2.5 h, 4°C. After centrifugation, gradients were fractionated using a Density Gradient Fraction System (Brandel, BR-188). To normalize for fraction volume in subsequent RT-qPCR experiments, 100 pg of *in vitro* transcribed RNA containing *Renilla* and *Firefly* luciferase (*in vitro* transcribed from the pRF-HCV vector) were added to each fraction. Equal volumes of fractions were then pooled as described in the text. RNA was then extracted by mixing samples with Acid-Phenol:Chloroform, pH 4.5 (with IAA, 125:24:1) (Thermo Fisher, AM9722), incubating at 65°C, 5 min, and subsequent centrifugation at 21,000xg, 10 min, RT. The aqueous phase was obtained, mixed 1:1 with 100% ethanol, and further purified using the RNA Clean & Concentrator-5 kit (Zymo Research, R1013). Samples were treated 30 minutes with TURBO DNase per manufacturer protocol (Thermo Fisher, AM2238), and subsequently purified again using the RNA Clean & Concentrator-5 kit. Next, 20 ng of RNA was used for first strand cDNA synthesis using the iScript Supermix (Bio-rad, 1708841). cDNA was diluted 20-fold and 4 µL was used in a reaction for SYBR green detection with SsoAdvanced SYBR Green supermix (Bio-Rad, catalog no. 1725270) on a CFX384 machine (Bio-rad). All primer sequences are available in Table S2.

For analysis, the candidate transcript qPCR Ct value in each pooled fraction was normalized to that of the spike-in *in vitro* transcribed *Renilla* luciferase RNA. Normalized Ct values were converted from log to linear space, multiplied by the number of fractions combined per pool, and then normalized to the sum total across all pooled fractions for a given sample. These values were compared between genotypes for each fraction using a paired, two-tailed t-tests with samples paired based on day of collection and somite count.

### Ribosome profiling of embryonic limb buds

Ribosome profiling was performed as described before(Ingolia et al., 2012) with modifications. Details are described below. E10.5 mouse embryonic limbs were harvested in cold HBSS (Thermo Fisher, 14025-076) containing 100 μg/ml cycloheximide (CHX). Choice of RNase was particularly important given that low input samples from a pair of E10.5 embryonic limbs were used. Although RNase I offers better codon resolution than RNase A/T1, RNase I has been shown to degrade ribosomes when used in high concentrations relative to input RNA whereas RNase A and T1 better maintain ribosome integrity at a greater range of concentrations despite only cutting single stranded RNA at C/U and G, respectively(Cenik et al., 2015). Given that E10.5 embryonic limbs yield low amounts of RNA, RNase A and T1 were used to prevent excess ribosome degradation.

Embryonic heads were collected for subsequent genotyping. Limb pairs from each embryo were lysed via vigorous pipetting and 30 minute incubation at 4°C in 215 μL buffer A (20 mM Tris-HCl pH 7.5, 150 mM NaCl, 15 mM MgCl2, 1 mM DTT, 8% glycerol, 1% Triton X-100, 100 μg/ml CHX (Sigma-Aldrich, C7698-1G), 20 U/ml Turbo DNAse (Thermo Fisher, AM2238), and Complete Protease Inhibitor EDTA-free (Sigma-Aldrich, 11836170001). Lysates were cleared via sequential centrifugation at 1,300g, 5 min and 10,000g, 10 min at 4°C. For RNA input for RNA-Seq, 70 μL of lysate was diluted in 55 μL of water and stored at -80°C in 375 μL of TRIzol LS (Thermo Fisher, 10296010). For ribosome profiling, 120 μL of cleared lysate was treated with 0.5 μg RNase A (Thermo Fisher, AM2271) and 300 U RNase T1 (Thermo Fisher, EN0541) for 30 min, RT with gentle rocking. The reaction was stopped by the addition of 100 U SUPERase RNase Inhibitor (Thermo Fisher, AM2694). Ribosomes were enriched by adding 110 μL lysate onto 900 μL sucrose cushion buffer (1 M sucrose in buffer A containing 20 U/mL SUPERase RNase Inhibitor), and centrifuging in a TLA 120.2 rotor (Beckman) 70,000 rpm, 4 h, 4 °C. The ribosome pellet containing the ribosome footprinted Ribo-Seq library was resuspended in 500 μL TRIzol.

Library preparation was adapted from previous protocols(Flynn et al., 2015; Ingolia et al., 2012) and the ARTseq Ribosome Profiling Kit manual (Epicentre, Illumina). In summary, total RNA and ribosome footprints were extracted using the Direct-zol Micro Kit (Zymo, R2060) with in-column DNase I treatment. To have an adequate amount of RNA for subsequent steps, pairs of samples of the same genotype were pooled to generate a single replicate sample, brought up to 90 μL with water, and precipitated with 150 μL isopropanol overnight -80°C after addition of 10 μL 3 M NaOAc pH 5.5 and 1.5 μL 15 mg/mL GlycoBlue Coprecipitant (Thermo Fisher, AM9515). The samples were then centrifuged at 21,000g, 30 min, 4°C, supernatant was removed, and the RNA pellet was washed 500 μL cold 75% ethanol. Pellets were dried for 10 min, RT and resuspended in 15 μL nuclease free water. After extraction and precipitation, both ribosome footprinting and total RNA samples were depleted of rRNA using the Ribo-Zero Gold rRNA Removal Kit (H/M/R) (Illumina, catalog no. MRZG126) with the modification that the 50°C incubation was not performed for the ribosome footprinting samples. The samples were then column purified (RNA Clean & Concentrator 5, Zymo Research, catalog no. R1016) with the modifications that 2 volumes of RNA Binding Buffer and 4.5 volumes of ethanol were added to ribosome footprinting samples to purify small RNAs and 1 volume of RNA Binding Buffer and 1 volume of ethanol was added to total RNA samples to isolate RNA > 200 nt. Total RNA samples were then fragmented by partial alkaline hydrolysis. The samples were diluted to 100 μL with 5 mM Tris-HCl, pH 7.5 and incubated with 100 μL 2x alkaline fragmentation buffer (100 mM Na2CO3 pH 9.2, 2 mM EDTA) for 20 minutes at 95°C. The reaction was neutralized with 440 μL STOP Buffer (70 μL 3 M NaOAc pH 5.5, 2 μL Glycoblue, and 370 μL nuclease free water) and isopropanol precipitated overnight at -80°C.

Ribosome protected fragments and total RNA samples were then size selected by running the samples out on a 15% TBE-Urea polyacrylamide gel. Ribosome protected fragments were size selected between 28-nt and 34-nt as marked by RNA oligonucleotides oNTI199 and oNTI265, respectively(Ingolia et al., 2011). Total RNA samples were size selected between 34-70 nt as marked by a 10 bp DNA ladder (Invitrogen, catalog no. 10821015). Gel slices were crushed and extracted at room temperature overnight in 400 μL RNA extraction buffer (300 mM NaOAc pH 5.5, 1 mM EDTA, 0.25% SDS) and isopropanol precipitated. Samples were then 3’dephosphorylated by denaturing at 65°C for 5 min and incubating with T4 PNK (NEB, catalog no. M0201S) in a 10 μL reaction (7 μL precipitated RNA, 1 μL 10x T4 PNK Buffer, 1 μL SUPERase Inhibitor, 1 μL 10 U/μL T4 PNK) at 37°C for 1 hour. The reaction was stopped through heat inactivation at 65°C for 20 min. To ligate adaptor, samples were then incubated with 0.5 μL of 50 μM Universal miRNA Cloning Linker (NEB, catalog no. S1315S) and denatured at 65°C for 5 min. The denatured sample was then incubated with 1 μL T4 RNA Ligase 2, truncated KQ (NEB, M0373S), 1 μL 10x buffer, 6 μL 50% PEG 8000, and 1.5 μL water for 4.5 h at 25°C. Free adaptor was then removed by addition of 1 μL 10 U/μL 5’-Deadenylase (NEB, M0331S). 1 μL 10 U/μL RecJ Exonuclease (Lucigen/Epicentre, RJ411250). and 1 μL 20 U/μL SUPERase Inhibitor. Samples were then purified using Zymo RNA Clean & Concentrator-5 columns using the protocol above to preserve small RNAs (100 μL sample, 200 μL RNA binding buffer, 450 μL 100% ethanol).

To perform reverse transcription, 10 μL sample was incubated with 2 μL of 1.25 μM RT primer (see Table S2) and denatured at 65°C for 5 min. Reverse transcription (RT) was performed with primers containing sample barcodes and unique molecular identifiers. RT was performed with SuperScript III (Thermo Fisher, 18080-044) in a 20 μL reaction (48°C, 40 min). RNA was then hydrolyzed by adding 2.2 μL of 1N NaOH and incubating for 20 min at 98°C. Samples were then purified using Zymo RNA Clean & Concentrator-5 columns (100 μL sample, 200 μL RNA binding buffer, 450 μL 100% ethanol). RT products were size selected and gel extracted from 10% TBE Urea polyacrylamide gels as described above instead that DNA extraction buffer was used for overnight extraction (300 mM NaCl, 10 mM Tris-HCl pH 8, 1 mM EDTA, 0.1% SDS). Eluate was then isopropanol precipitated overnight at -80°C overnight.

Samples were then circularized with CircLigase (Illumina, CL4115K) in a 20 μL reaction (15 μL cDNA, 2 μL 10x CircLigase Buffer, 1 μL 1 mM ATP, 1 μL MnCl2, 1 μL CircLigase) for 12 hours at 60°C and subsequently purified by Zymo RNA Clean & Concentrator 5 columns (100 μL sample, 200 μL RNA binding buffer, 450 μL 100% ethanol) and eluted with 12 μL 10 mM Tris-HCl pH 8. 1 μL of library was used for PCR amplification with Phusion High-Fidelity DNA Polymerase (Thermo Fisher, catalog no. F530S) (98°C 30s, 98°C 10s, 65°C 10s, 72°C 5s) for 9-10 cycles (see Table S2 for primers). PCR product was PAGE purified from native 8% TBE polyacrylamide gels, extracted overnight using DNA extraction buffer, and isopropanol precipitated for > 2 h at -80°C. DNA was measured and quality controlled on the Agilent 2100 Bioanalyzer (High-Sensitivity DNA) by the Stanford Protein and Nucleic Acid (PAN) Facility. Libraries were sequenced by the Stanford Functional Genomics Facility (SFGF) on the Illumina NextSeq 500 (1x75nt).

### Ribosome profiling analysis

For analysis pre-processing, UMI-tools(Smith et al., 2017) was used to extract sample barcodes and UMIs. Reads were demultiplexed, sample barcodes were removed, and reads were quality filtered (fastq_quality_filter -Q33 -q 20 -p 70) using FASTX-Toolkit (http://hannonlab.cshl.edu/fastx_toolkit/). The 3’ adapter sequence CTGTAGGCACCATCAAT was removed using cutadapt(Martin, 2011), and the 5’ end of each read was then removed using fastx_trimmer from FASTX-Toolkit. To remove reads that aligned to rRNA, tRNA, and snRNAs, reads were first aligned to these sequences using bowtie2(Langmead and Salzberg, 2012) (bowtie2 parameters: -L 18) and subsequently discarded. Filtered reads that did not align to rRNA/tRNA/snRNAs were then aligned using bowtie2 (bowtie2 parameters: --norc -L 18) to an GRCm38/mm10 transcriptome reference derived from UCSC/GENCODE VM20 knownCanonical annotations filtered for high confidence transcripts that contain at least one of the following: a Uniprot ID, a RefSeq ID, or an Entrez ID. PCR duplicates were then removed using UMI-tools. Ribosome protected fragments (RPFs) were then parsed for uniquely aligned reads, separated into read length groups, and ribosome A site positions were determined by offsetting the distance of the 5’ end of each read to canonical start sites in each length group and adding 4 nucleotides(Ingolia et al., 2011). RNA-Seq reads were also parsed for uniquely aligned reads and were assigned to the center of each read. RPFs and RNA-Seq reads were then counted using the transcriptome annotations detailed above. Reads aligning to the CDS (with the first 15 codons and last 5 codons removed) were used for RPF libraries, and reads aligning to the entire transcript were used for RNA-Seq libraries. Transcripts with counts per million (cpm) > 0.75 for at least 3 ribosome profiling libraries were retained for downstream analysis. RPF and RNA-Seq libraries were then normalized separately by the method of trimmed mean of M-values (TMM) using edgeR(Robinson et al., 2010).

Differential translation analysis was performed using voom(Law et al., 2014) and limma(Ritchie et al., 2015). The following design matrix was constructed (∼0 + genotype and type of library (RPF vs. RNA) + replicate batch + sex). The corresponding contrast matrix comparing translational efficiency (RPF - RNA) for each of the pairs of genotypes of interest (e.g., *Prx1^Cre^;Rps6^lox/+^;Trp53^+/+^* - *Prx1^Cre^;Rps6^lox/+^;Trp53^+/+^*, *Prx1^Cre^;Rps6^lox/+^;Trp53^-/-^* - *Prx1^Cre^;Rps6^lox/+^;Trp53^-/-^*, etc.) was also constructed. The data was then voom transformed to remove mean-variance count heteroscedascity. Using the design and contrast matrices mentioned above, linear models were fit to the data using limma with empirical Bayes moderation. Significant differentially translated genes were defined as those with FDR < 0.1 obtained using the Benjamini-Hochberg method.

Gene set enrichment analysis was performed using Camera (Wu and Smyth, 2012) with mouse GO Biological Processes gene sets obtained from http://download.baderlab.org/EM_Genesets/current_release/Mouse/Entrezgene/. Gene sets were filtered such that all genes in gene sets have expression values in the dataset. Gene sets were also filtered to only include those with > 10 and < 500 genes. Enriched gene sets were visualized using Enrichment Map (Merico et al., 2010) and Cytoscape. For feature analyses, UTR RNA folding energies were obtained from UCSC Genome Browser (foldUtr5 and foldUtr3).

To select for high-confidence translationally repressed transcripts involved in limb development, transcripts were selected such that 1) the TE was significantly decreased in *Prx1^Cre^;Rps6^lox/+^* vs. *Rps6^lox/+^* embryos, and 2) the transcripts also had a significant decrease in RPF, which filters for transcripts whose changes in TE are driven primarily by ribosome footprints and not by changes in RNA transcript levels. The transcripts that then intersected with the significantly enriched limb-related GO categories (appendage development, appendage morphogenesis, limb development, and limb morphogenesis) were selected.

### Samples and Statistical Analysis

All measurements were sampled from individual biological replicates. Replicates for mouse embryo experiments consisted of individual limbs, individual embryos, or pools of embryos if material was limiting. Experiments consisted of multiple embryos from multiple litters. Unless otherwise stated, when groups of continuous data are compared, two-tailed t-tests with unequal variance were performed. No statistical methods were used to predetermine sample size.

### Code Availability

All custom codes are available upon request from the corresponding author.

### Data Availability

Data that support the findings of this study have been deposited in the Gene Expression Omnibus under accession number GSE135722.

## SUPPLEMENTARY TABLES

**Table S1.** Limb phenotype quantification. Related to Figures 1 and S1.

**Table S2.** Primers and oligos.

**Table S3.** Ribosome profiling results from *Prx1^Cre^;Rps6^lox/+^* vs. *Rps6^lox/+^* E10.5 limbs. Related to Figures 4 and S10.

**Table S4.** Ribosome profiling results from *Prx1^Cre^;Rps6^lox/+^;Trp53^-/-^* vs. *Rps6^lox/+^;Trp53^-/-^* E10.5 limbs. Related to Figures 4 and S10.

**Table S5.** Gene set enrichment analysis of translational efficiency changes in *Prx1^Cre^;Rps6^lox/+^* vs. *Rps6^lox/+^* E10.5 limbs. Related to Figure 4.

**Table S6.** RNA folding energies and lengths of transcripts. Related to Figure 4.

## SUPPLEMENTAL FIGURES

**Figure S1.**
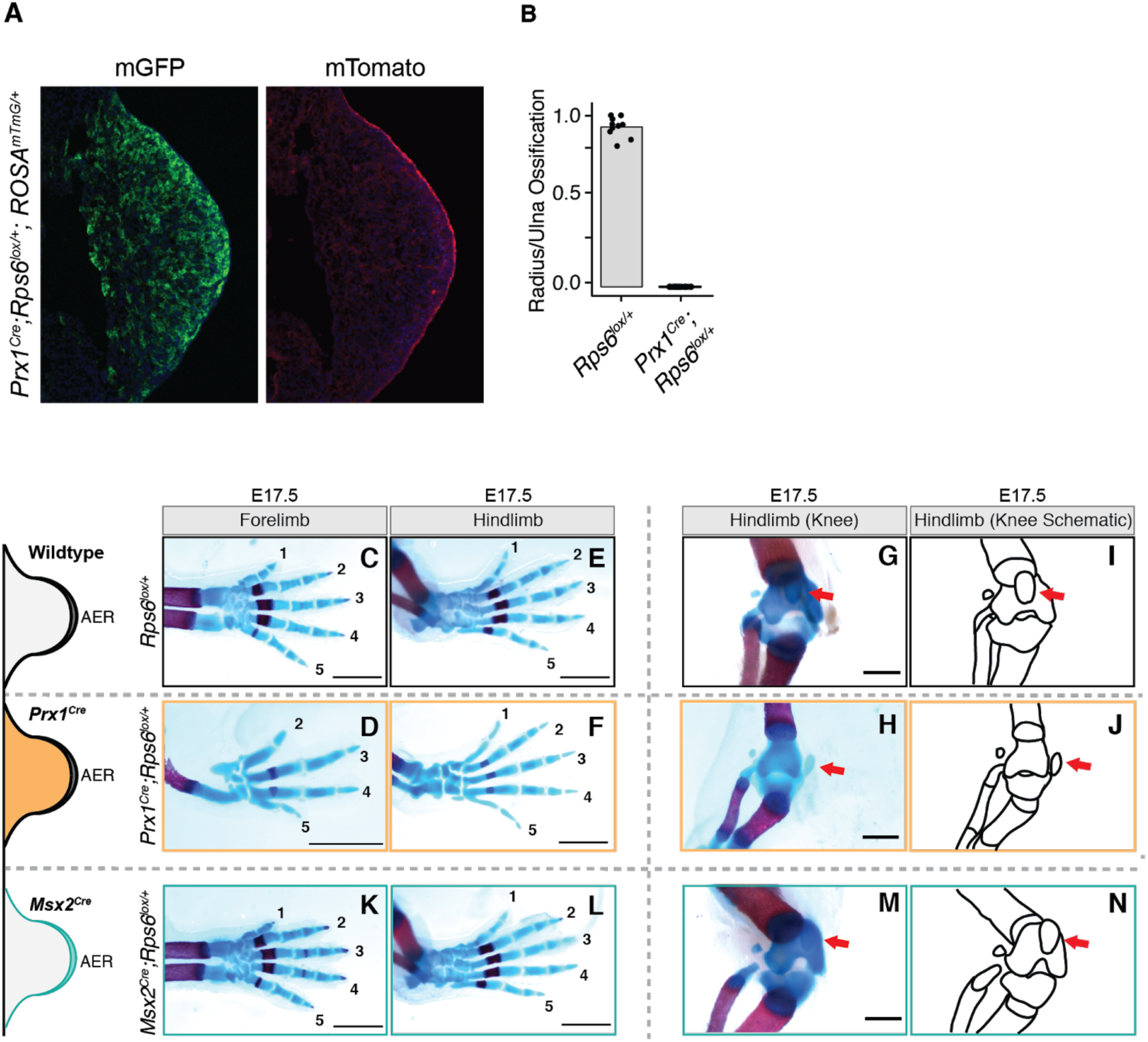
Conditional deletion of one *Rps6* allele leads to patterning defects. Related to Figure 1. (**A**) Representative tissue section from E9.5 *Prx1^Cre^;Rps6^lox/+^;Rosa^mTmG/+^* forelimb bud demonstrating distribution of recombination in the developing limb bud. green = recombined membrane GFP; red = unrecombined membrane Tomato. (**B**) Radius ossification normalized to ulna ossification in E17.5 forelimbs. *n* = 10 limbs (*Rps6^lox/+^*), *n* = 28 limbs (*Prx1^Cre^;Rps6^lox/+^*). (**C-N**) Characterization of phenotypes associated with *Rps6* haploinsufficiency. **Left**: Digit phenotypes found in the forelimb and hindlimb of wildtype *Rps6^lox/+^* (top row), *Prx1^Cre^;Rps6^lox/+^* (middle row), and *Msx2^Cre^;Rps6^lox/+^* (bottom row) E17.5 embryos, Numbers label digits. **Right**: Patella phenotypes found in the hindlimb of wildtype *Rps6^lox/+^* (top row), *Prx1^Cre^;Rps6^lox/+^* (middle row), and *Msx2^Cre^;Rps6^lox/+^* (bottom row) E17.5 embryos. Red arrows denote the patella.

**Figure S2.**
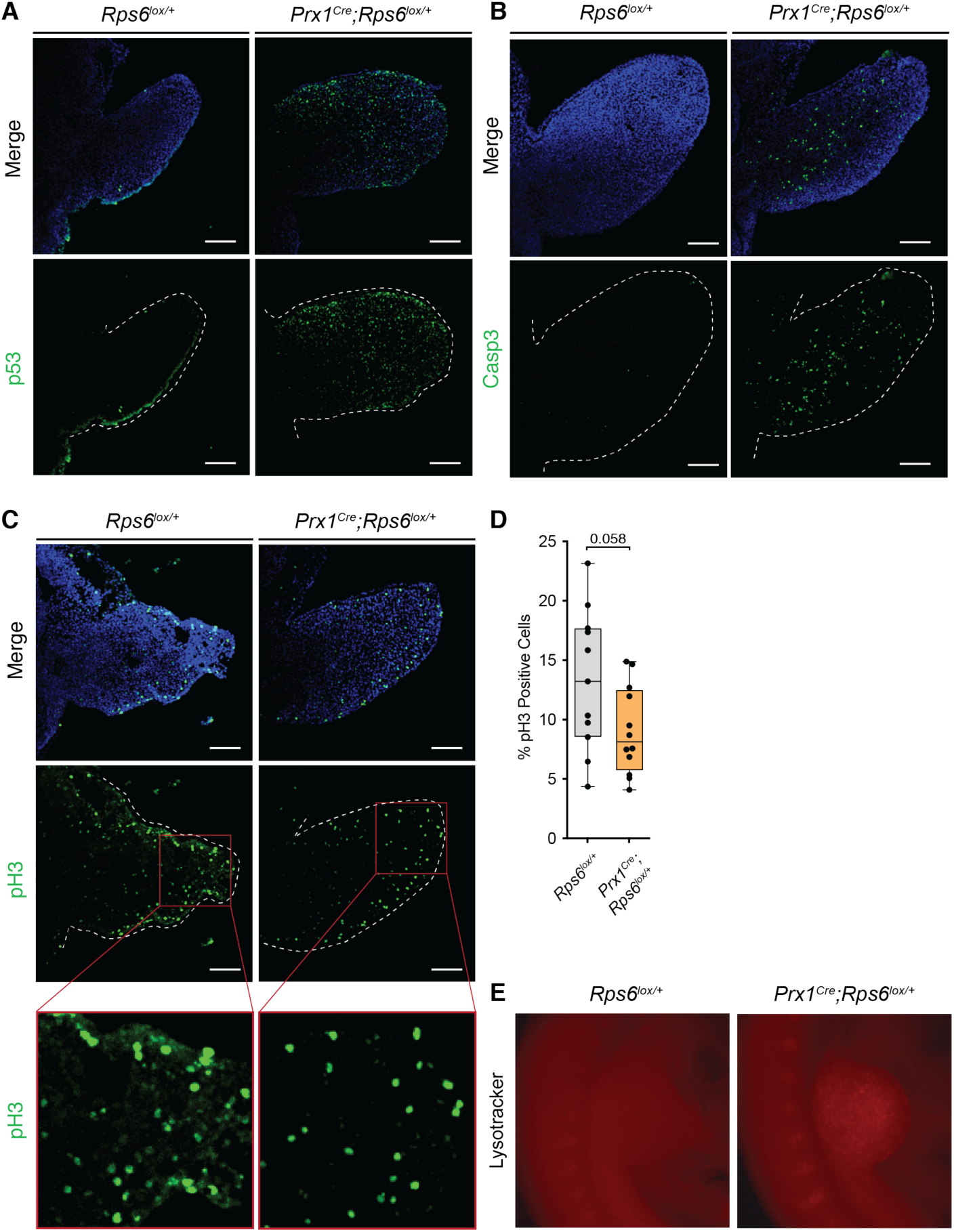
p53 activation is not spatially restricted in the developing forelimb. Related to Figure 1. (**A**) Tissue sections from E10.5 wildtype (*Rps6^lox/+^*) and *Prx1^Cre^;Rps6^lox/+^* forelimbs staining for p53 (green). DAPI staining shown in blue. (**B**) Tissue sections from E10.5 wildtype (*Rps6^lox/+^*) and *Prx1^Cre^;Rps6^lox/+^* forelimbs staining for cleaved Caspase-3 (green) to indicate apoptotic cells. DAPI staining shown in blue. (**C**) Tissue sections from E10.5 wildtype (*Rps6^lox/+^*) and *Prx1^Cre^;Rps6^lox/+^* forelimbs staining pH3 (green) to indicate actively proliferating cells. DAPI staining shown in blue. (**D**) Percentage of pH3-labeled cells from tissue sections of E10.5 wildtype (*Rps6^lox/+^*) and *Prx1^Cre^;Rps6^lox/+^* forelimbs. *n* = 11 sections from 3 embryos wildtype, *n* = 12 sections from 3 embryos *Prx1^Cre^;Rps6^lox/+^*. For box-plots, center line, median; box limits, first and third quartiles; whiskers, 1.5x interquartile range; points, outliers, * *P* < 0.05, two-tailed t-test, unequal variance. (**E**) Lysotracker Red stained E10.5 forelimbs from wildtype (*Rps6^lox/+^*) and *Prx1^Cre^;Rps6^lox/+^* embryos as a marker of cell death.

**Figure S3.**
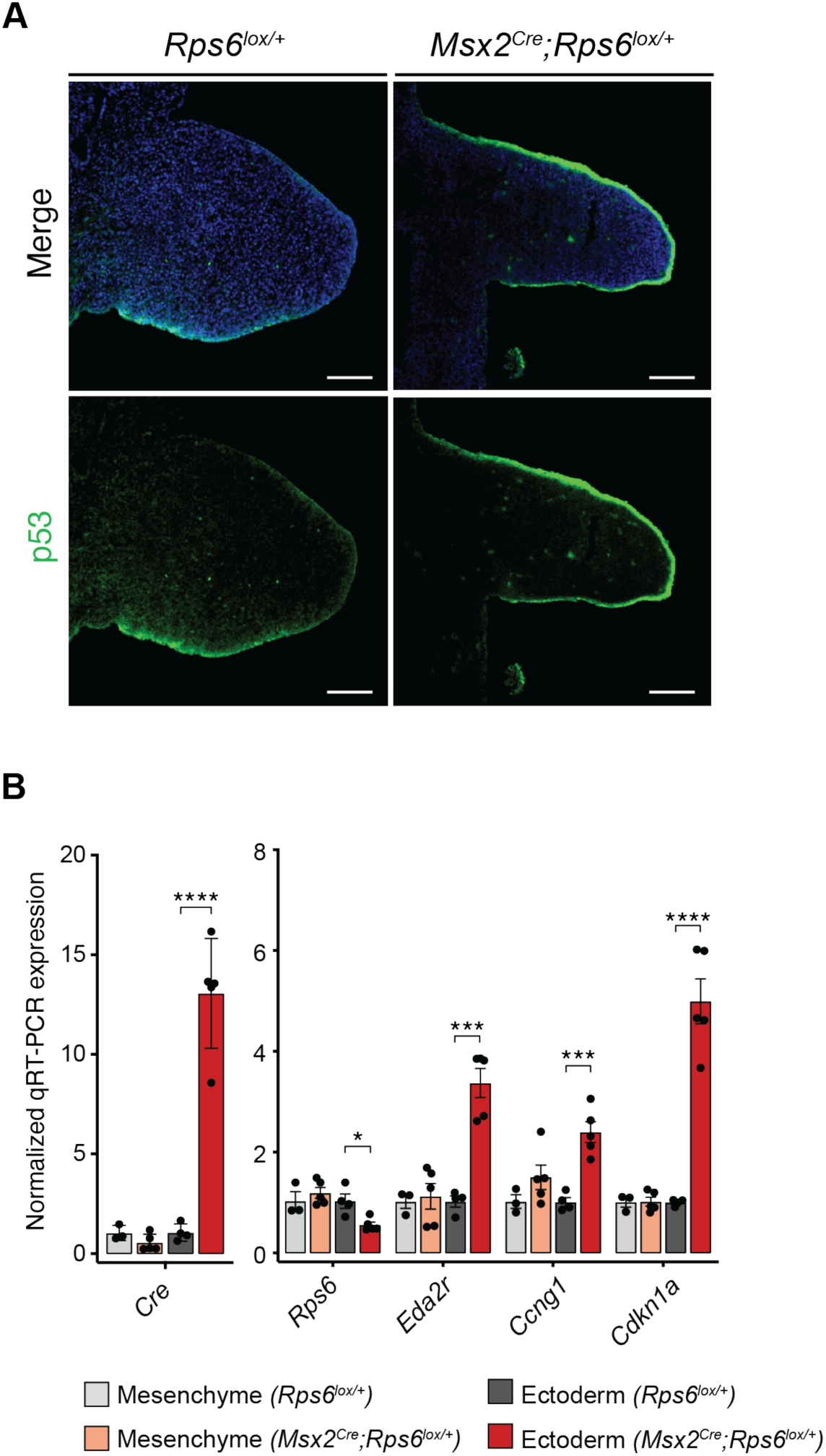
p53 is active in the AER of *Msx2^cre^;Rps6^lox/+^* limbs. Related to Figure 1. (**A**) Tissue sections from E10.5 wildtype (*Rps6^lox/+^*) and *Msx2^Cre^;Rps6^lox/+^* forelimbs staining for p53 (green) and DAPI (blue). (**B**) RT-qPCR of select p53 target genes and *Rps6* mRNA from isolated ectoderm or mesenchyme layers from E11.5 wildtype (*Rps6^lox/+^*) and *Msx2^Cre^;Rps6^lox/+^* forelimbs. Expression was normalized to *Actb* and *NupL1* as housekeeping genes and then to the mean of wildtype for each tissue. Error bars = SEM, * *P* < 0.05, ** *P* < 0.01, *** *P* < 0.001, **** *P* < 0.0001, two-tailed t-test, unequal variance.

**Figure S4.**
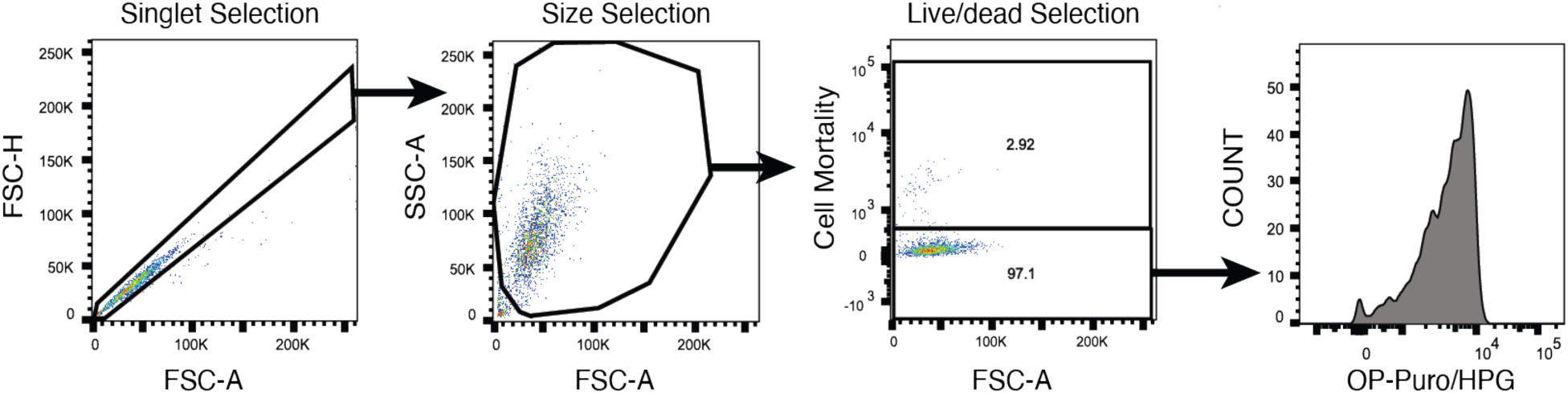
Overview of flow cytometry gating schemes for OP-Puro and HPG labeling experiments. Related to Figure 1. Gating scheme for OP-Puro and HPG incorporation assays. FACS gates of analyzed cells labeled with Zombie Violet Live-Dead Stain and Alexa Fluor 555 Picolyl Azide dye after incorporation of OP-puro or HPG.

**Figure S5.**
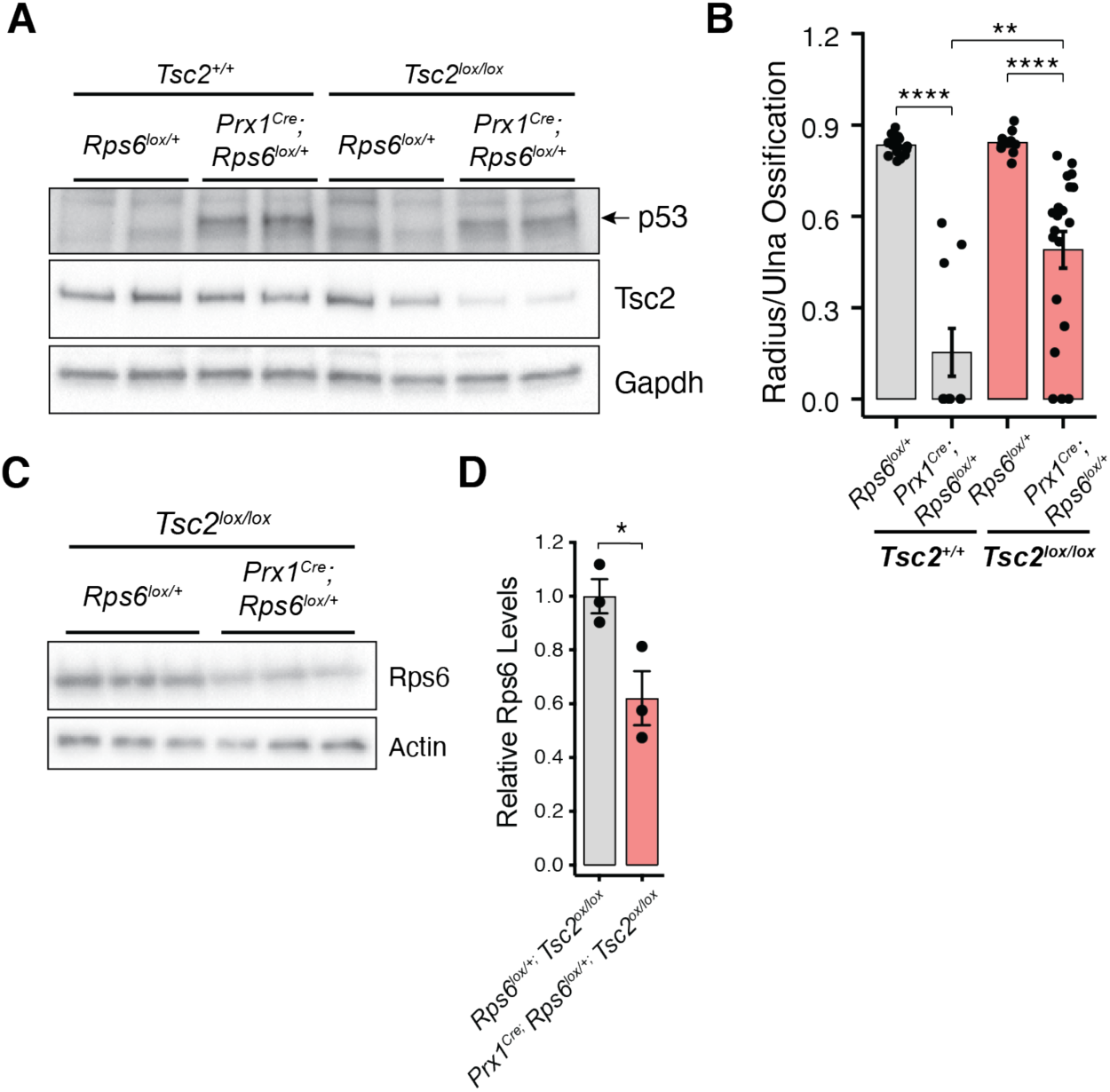
Loss of *Tsc2* rescues the *Rps6* haploinsufficient limb phenotype and increases translation in the presence of p53 activation. Related to Figure 2. (**A**) Western blot of p53 and Tsc2 from developing E10.5 forelimbs of *Rps6^lox/+^;Tsc2^+/+^*, *Prx1^Cre^;Rps6^lox/+^;Tsc2^+/+^*, *Rps6^lox/+^;Tsc2^lox/lox^*, and *Prx1^Cre^;Rps6^lox/+^;Tsc2^lox/lox^* embryos showing that p53 is still activated upon *Rps6* haploinsufficiency in a *Tsc2* conditional null background. (**B**) Quantification of radius/ulna ossification in E17.5 forelimbs. *n* = 18 limbs, (*Rps6^lox/+^;Tsc2^+/+^*); *n* = 10 limbs, (*Prx1^Cre^;Rps6^lox/+^;Tsc2^+/+^*); *n* = 10 limbs, (*Rps6^lox/+^;Tsc2^lox/lox^*); *n* = 20 limbs, (*Prx1^Cre^;Rps6^lox/+^;Tsc2^lox/lox^*). (**C**) Western blot of Rps6 from E10.5 forelimbs of *Rps6^lox/+^;Tsc2^lox/lox^* and *Prx1^Cre^;Rps6^lox/+^;Tsc2^lox/lox^* embryos showing that Rps6 is still decreased upon *Rps6* haploinsufficiency in a *Tsc2* conditional null background. (**D**) Quantification of Rps6 levels in (C) normalized to wildtype embryos. Error bars = SEM, * *P* < 0.05, ** *P* < 0.01, *** *P* < 0.001, **** *P* < 0.0001, two-tailed t-test, unequal variance.

**Figure S6.**
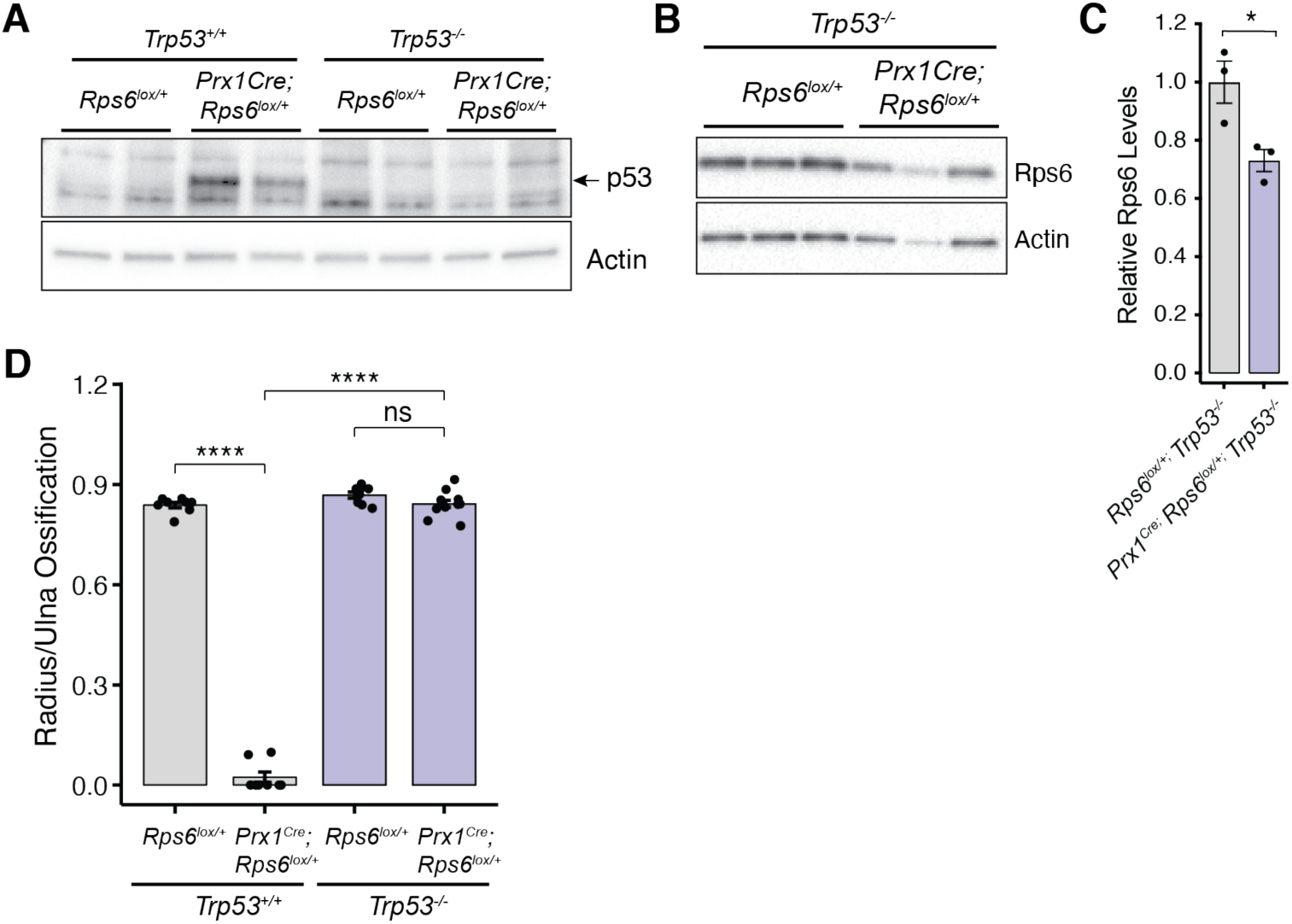
p53 deletion rescues radius phenotype and translation levels upon *Rps6* haploinsufficiency. Related to Figure 3. (**A**) Western blot of p53 from E10.5 forelimbs from *Rps6^lox/+^;Trp53^+/+^*, *Prx1^Cre^;Rps6^lox/+^;Trp53^+/+^*, *Rps6^lox/+^;Trp53^-/-^*, and *Prx1^Cre^;Rps6^lox/+^;Trp53^-/-^* embryos. (**B**) Representative Western blot of RPS6 from *Rps6^lox/+^;Trp53^-/-^*, and *Prx1^Cre^;Rps6^lox/+^;Trp53^-/-^* forelimbs at E10.5 showing that Rps6 is still decreased upon *Rps6* haploinsufficiency in a *Trp53* null background. (**C**) Quantification of Rps6 levels in (B) normalized to wildtype embryos. (**D**) Quantification of radius/ulna ossification in E17.5 forelimbs. *n* = 8 limbs, (*Rps6^lox/+^;Trp53^+/+^*); *n* = 8 limbs, (*Prx1^Cre^;Rps6^lox/+^;Trp53^+/+^*); *n* = 8 limbs, (*Rps6^lox/+^;Trp53^-/-^*); *n* = 12 limbs, (*Prx1^Cre^;Rps6^lox/+^;Trp53^-/-^*). Error bars = SEM, * *P* < 0.05, ** *P* < 0.01, *** *P* < 0.001, **** *P* < 0.0001, two-tailed t-test, unequal variance.

**Figure S7.**
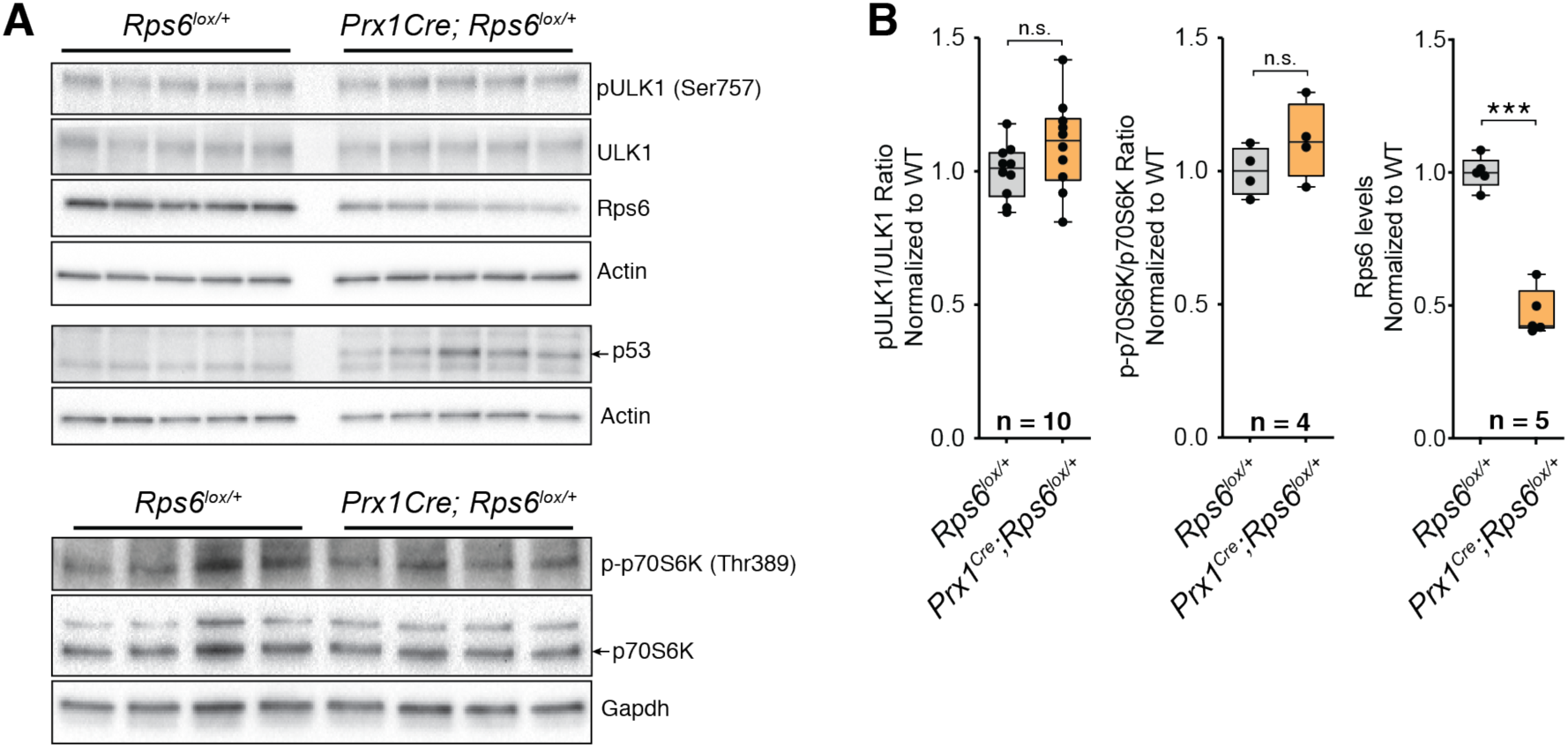
*Rps6* haploinsufficiency demonstrates an increase in p53 protein, a decrease in Rps6, and no significant perturbation of mTORC1 pathway activity. Related to Figures 1-3. (**A**) Representative Western blots of p53, Rps6, and phosphorylated proteins targeted by the mTORC1 pathway (ULK1, p70S6K). Each lane represents forelimbs from an individual *Rps6^lox/+^* or *Prx1^Cre^;Rps6^lox/+^* E10.5 embryo. (**B**) Quantification of Western blots for Rps6 in addition to mTOR pathway targets. For mTOR pathway targets, each protein was normalized to its non-phosphorylated form and then to wildtype *Rps6^lox/+^*. For phospho-ULK1, quantification was done for 10 limbs over two blots. Rps6 quantification was normalized to Gapdh and wildtype *Rps6^lox/+^*. For all blots equal amounts of limb lysate was used for detection. For box-plots, center line, median; box limits, first and third quartiles; whiskers, 1.5x interquartile range; points, outliers. * *P* < 0.05, two-tailed t-test, unequal variance.

**Figure S8.**
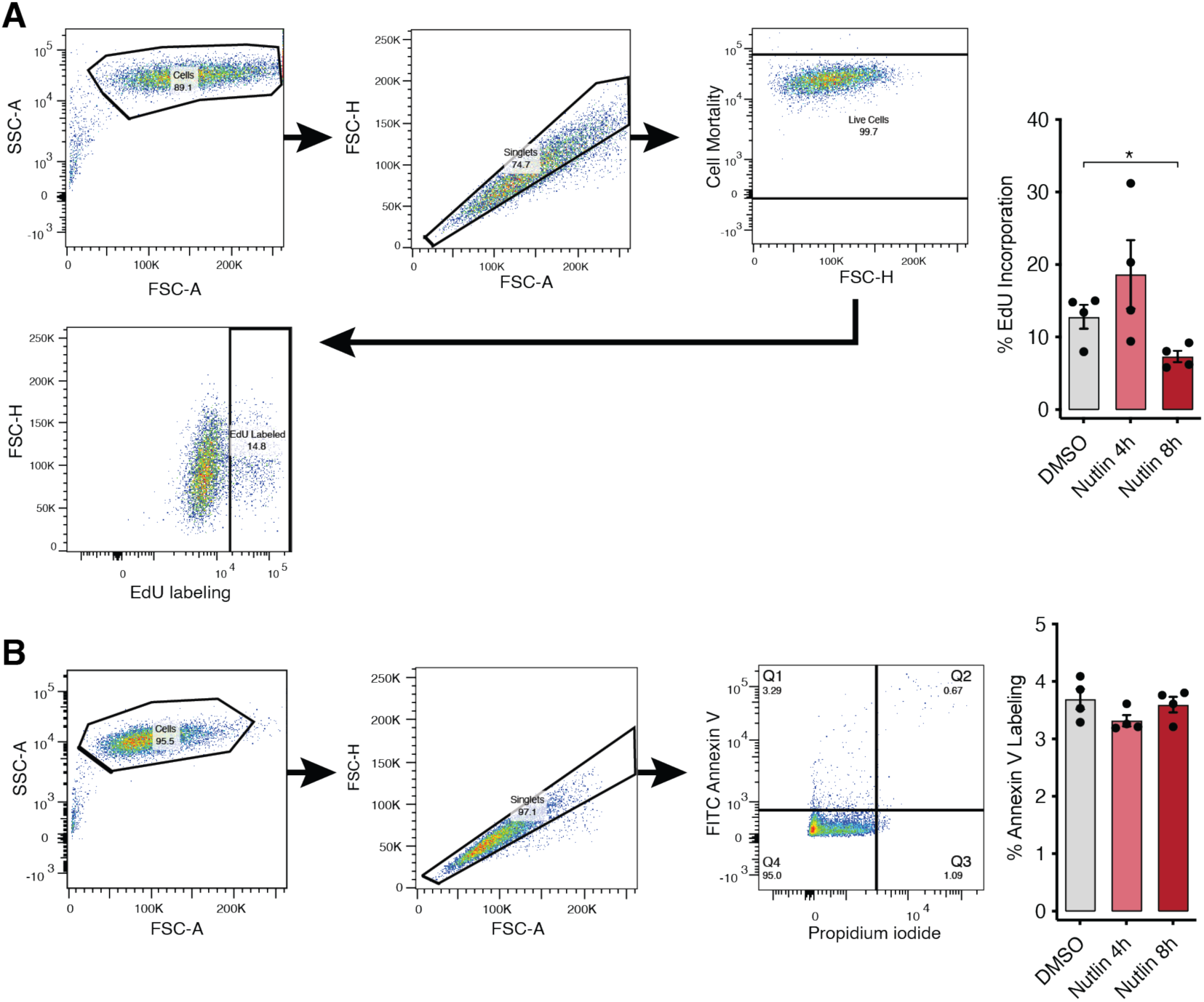
Nutlin-3a treatment causes translational repression with minimal effects on cell proliferation and apoptosis. Related to Figure 3. (**A**) FACS gating scheme of NIH3T3 fibroblast cells labeled with Zombie Violet Live-Dead Stain and Alexa Fluor 488 Azide dye after incorporation of EdU to measure cell proliferation. Right: percentage of EdU-positive cells after treatment with DMSO or Nutlin-3a for 4 and 8 h. (**B**) FACS gating scheme of NIH3T3 fibroblast cells stained with Annexin V and propidium iodide to measure levels of apoptosis. Right: percentage of Annexin V-labeled cells in Q1 after treatment with DMSO or Nutlin-3a for 4 and 8 h. Error bars = SEM, * *P* < 0.05, ** *P* < 0.01, *** *P* < 0.001, **** *P* < 0.0001, two-tailed t-test, unequal variance.

**Figure S9.**
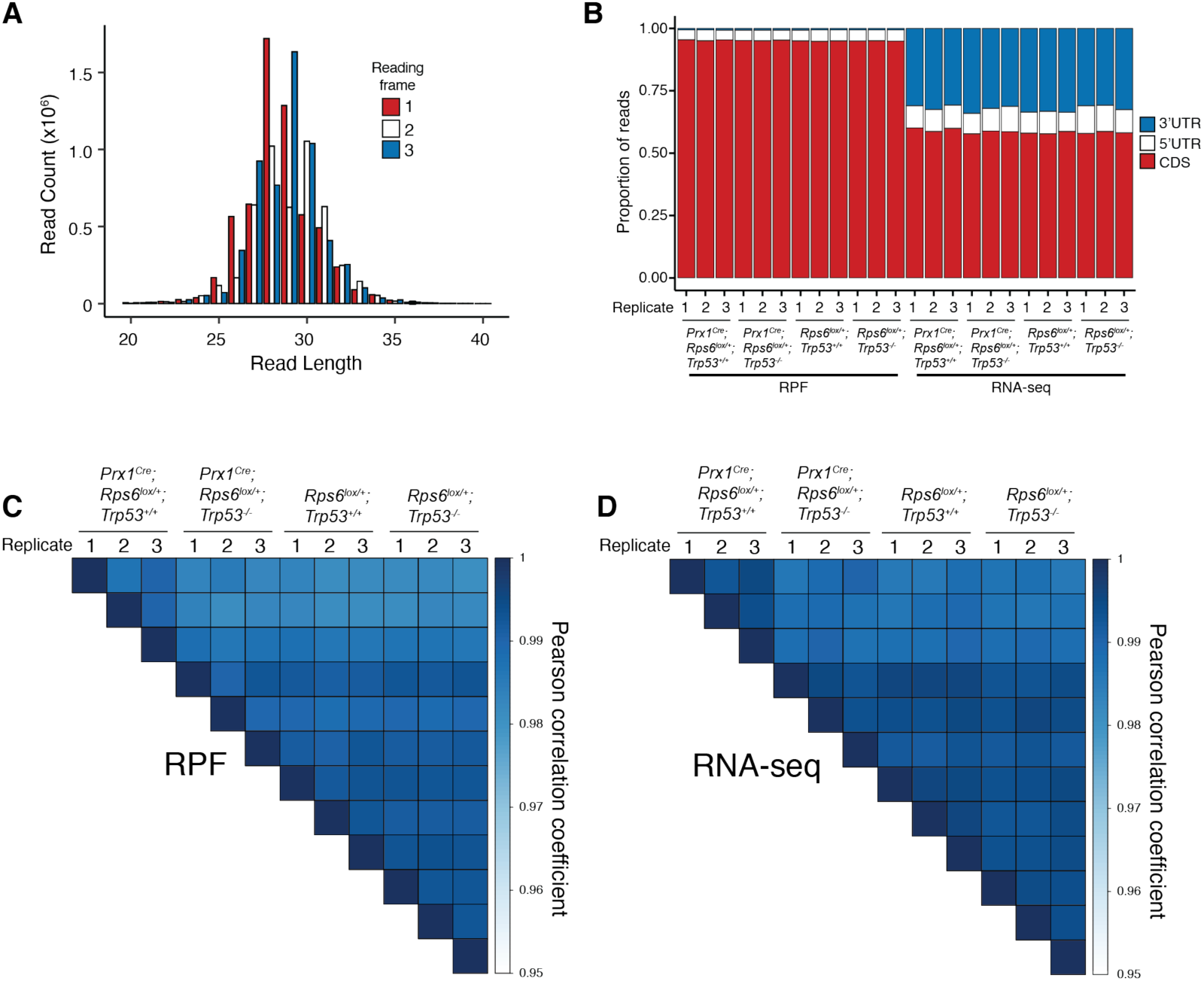
Quality control plots of ribosome profiling experiments demonstrate reading frame and CDS enrichment for ribosome protected fragments and good replicate reproducibility. Related to Figure 4. (**A**) Read length and CDS reading frame distribution for ribosome protected fragments (RPF) aligning to annotated ORFs in a representative library. Note that RNase A and T1 cut single-stranded RNA at only C/U and G, respectively, thus accounting for the slightly incoherent frame distribution (see Methods). (**B**) Proportion of reads aligning to CDS, 5’UTR, and 3’UTR regions from RPF and RNA-Seq libraries. Note the expected CDS enrichment for ribosome protected fragments. (**C-D**) Correlation matrix displaying sample-to-sample Pearson correlation coefficients for (C) RPF and (**D**) RNA-Seq log2(counts per million reads).

**Figure S10.**
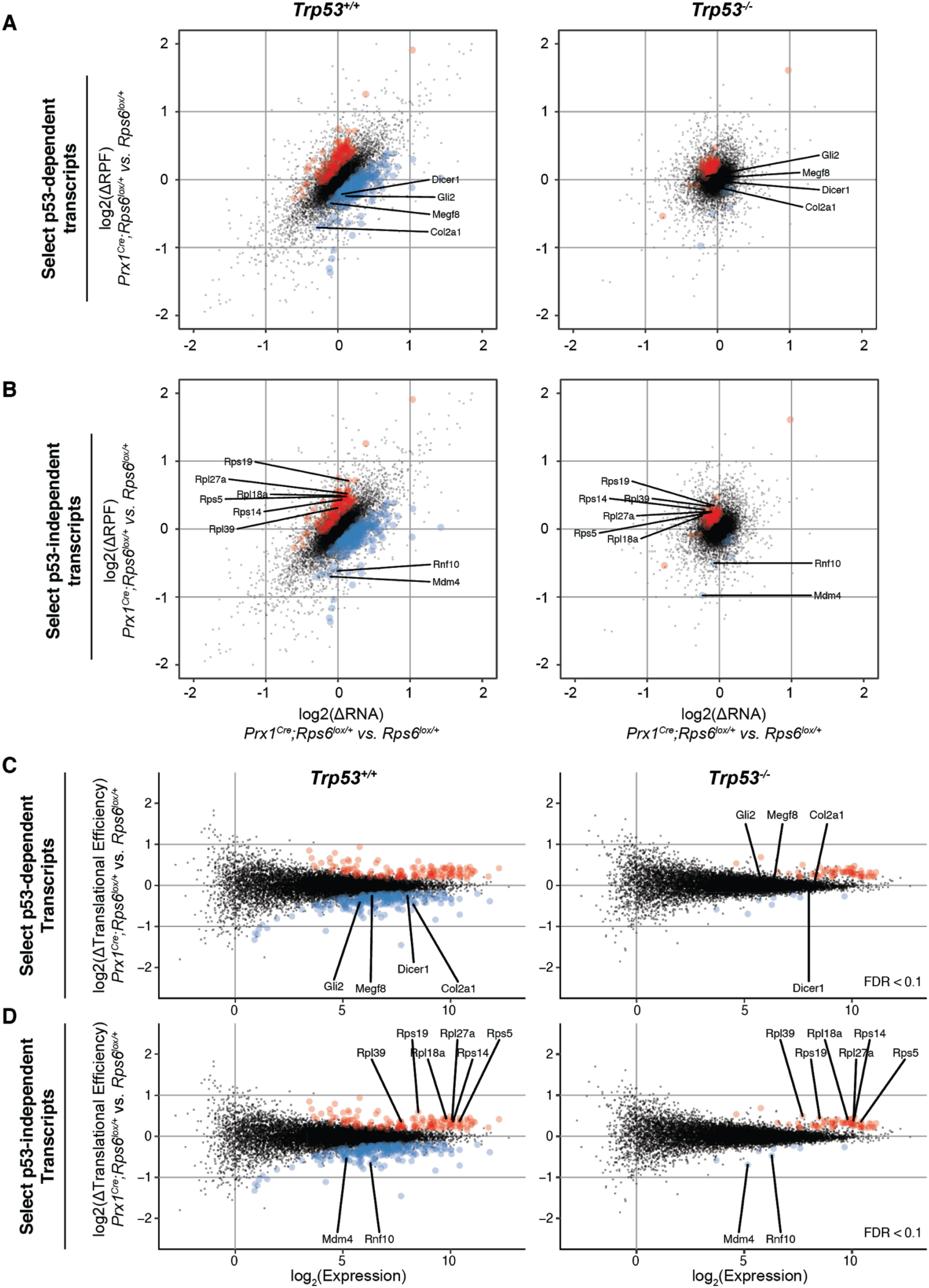
Analysis of ribosome profiling experiments. Related to Figure 4. (**A-B**) Scatter plots of changes in ribosome footprints (RPF) vs. RNA in E10.5 *Prx1^Cre^;Rps6^lox/+^* vs. *Rps6^lox/+^* embryonic forelimbs in a *Trp53^+/+^* or *Trp53^-/-^* background, highlighting select limb developmental transcripts with p53-dependent changes (A) and select transcripts with p53-independent changes (B). (**C-D**) Corresponding MA plots of change in translational efficiency (ΔTE) in E10.5 *Prx1^Cre^;Rps6^lox/+^* vs. *Rps6^lox/+^* embryonic forelimbs in a *Trp53^+/+^* or *Trp53^-/-^* background, highlighting select limb developmental transcripts with p53-dependent changes (C) and select transcripts with p53-independent changes (D). Note that 50 of the 64 significant translationally upregulated p53-independent transcripts consist of RPs (Table S4). Red, ΔTE > 0; blue, ΔTE < 0; *n* = 3 biological replicates (2 embryos each); FDR < 0.1.

**Figure S11.**
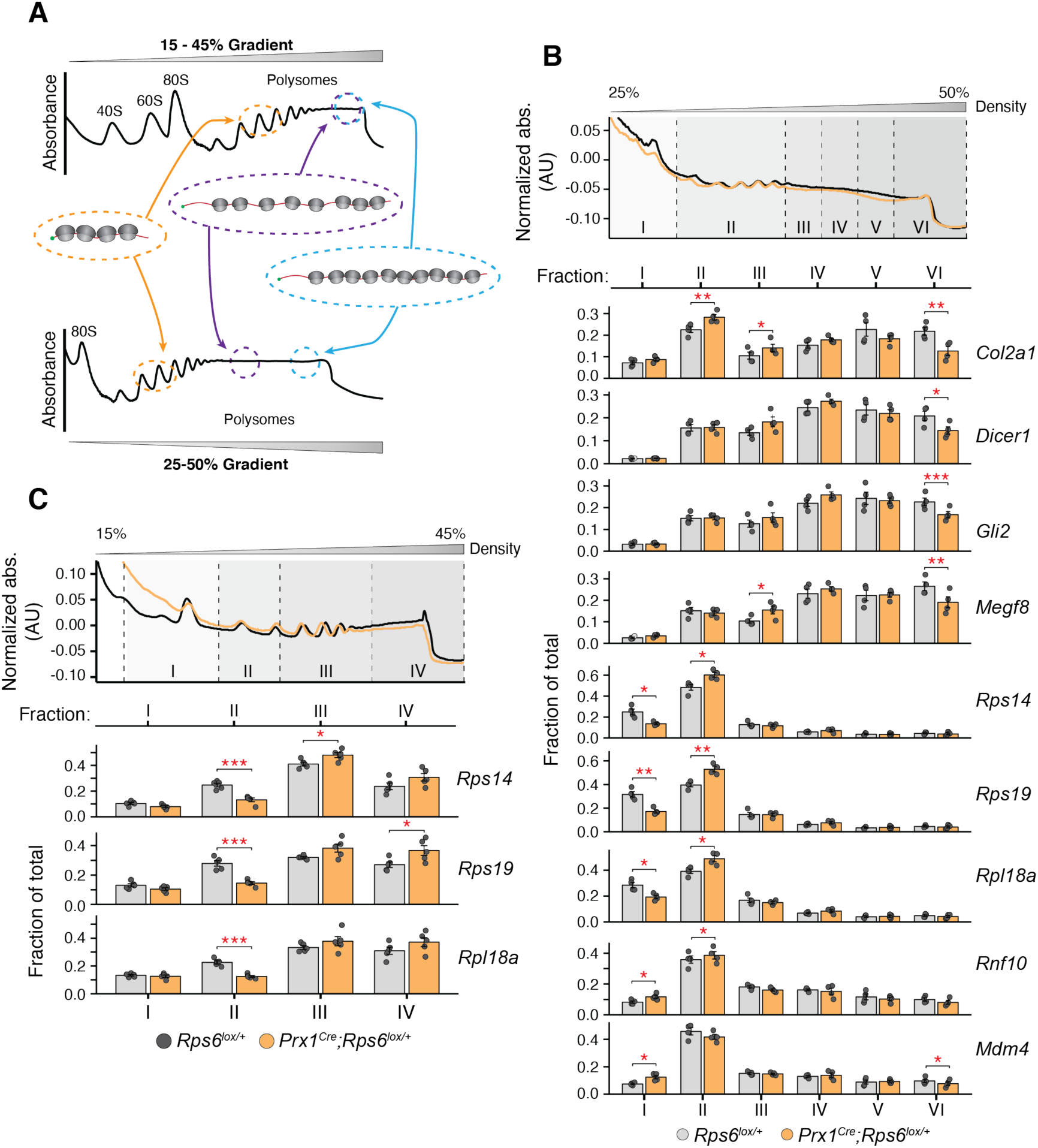
Sucrose gradient polysome fractionation and RT-qPCR validation of high-confidence differentially translated ribosome profiling candidate transcripts. Related to Figure 4. (**A**) Diagram of mRNAs co-fractionating with polysomes in sucrose density gradients. Note that mRNA transcript and ORF length impact the sensitivity of polysome gradients to resolve shifts in polysome profiles. Transcripts that are shorter in length inherently have less ribosomes bound, and thus appear earlier in the gradient fractions, while longer mRNAs naturally appear in the denser fractions of the gradient. Less dense gradients, which better separate transcripts bound to relatively few ribosomes, are therefore better at resolving changes in ribosome occupancy for shorter transcripts whereas more dense gradients, which better separate transcripts bound to many ribosomes, are better at resolving changes for longer transcripts. Given the strong length dependency of translationally changing transcripts upon *Rps6* haploinsufficiency (Figure 4E), different density gradients (15-45%, 25-50%) were performed to resolve changes in various length transcripts. (**B**) RT-qPCR of select mRNAs from 25-50% sucrose density gradients from E10.5 wildtype (grey) or *Prx1^Cre^;Rps6^lox/+^* (orange) limb buds. RT-qPCR Ct values were normalized to spike-in *in vitro* transcribed Renilla luciferase RNA added to each fraction and then to the sum total of all fractions. Top: polysome trace from wildtype (grey) or *Prx1^Cre^;Rps6^lox/+^* (orange) limb buds. Bottom: RT-qPCR of select mRNAs. *n =* 4 biological replicates. (**C**) RT-qPCR of select mRNAs from 15-45% sucrose density gradients from E10.5 wildtype (grey) or *Prx1^Cre^;Rps6^lox/+^* (orange) limb buds. RT-qPCR Ct values were normalized to spike-in *in vitro* transcribed Renilla luciferase RNA added to each fraction and then to the sum total of all fractions. Top: polysome trace from wildtype (grey) or *Prx1^Cre^;Rps6^lox/+^* (orange) limb buds. Bottom: RT-qPCR of select mRNAs. *n =* 5 biological replicates. * *P* < 0.05, ** *P* < 0.01, *** *P* < 0.001, two-tailed t-test, paired.

**Figure S12.**
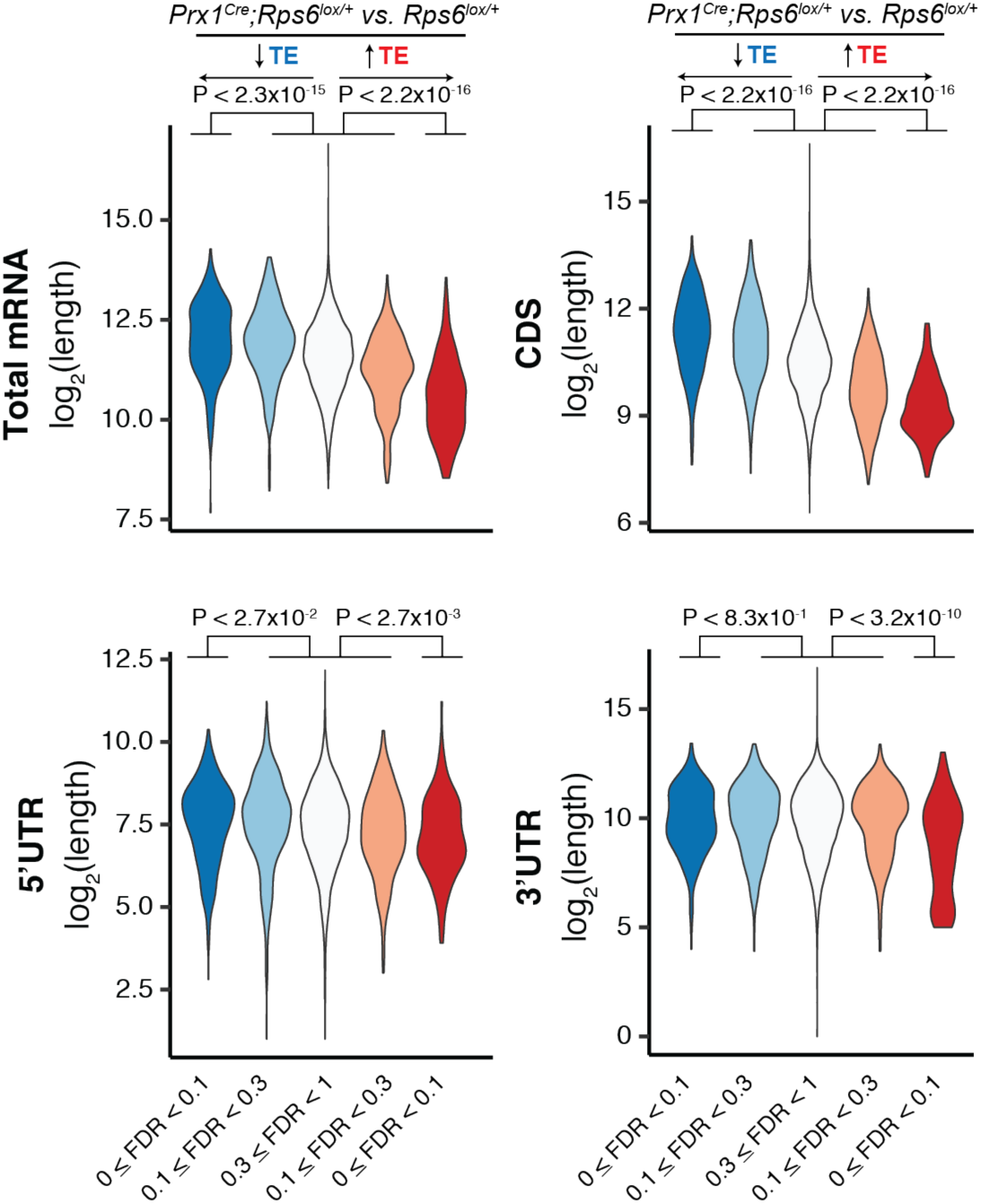
Transcript length, driven largely by CDS length, is negatively correlated with translation efficiency changes upon *Rps6* haploinsufficiency. Related to Figure 4. Violin plots quantifying ΔTE relative to total mRNA, CDS, 5’UTR, and 3’UTR length. Transcripts are stratified by direction of TE change (blue, down; red, up) and FDR (0 ≤ FDR < 0.1, 0.1 ≤ FDR < 0.3, 0.3 ≤ FDR ≤ 1); Mann-Whitney U test.

**Figure S13.**
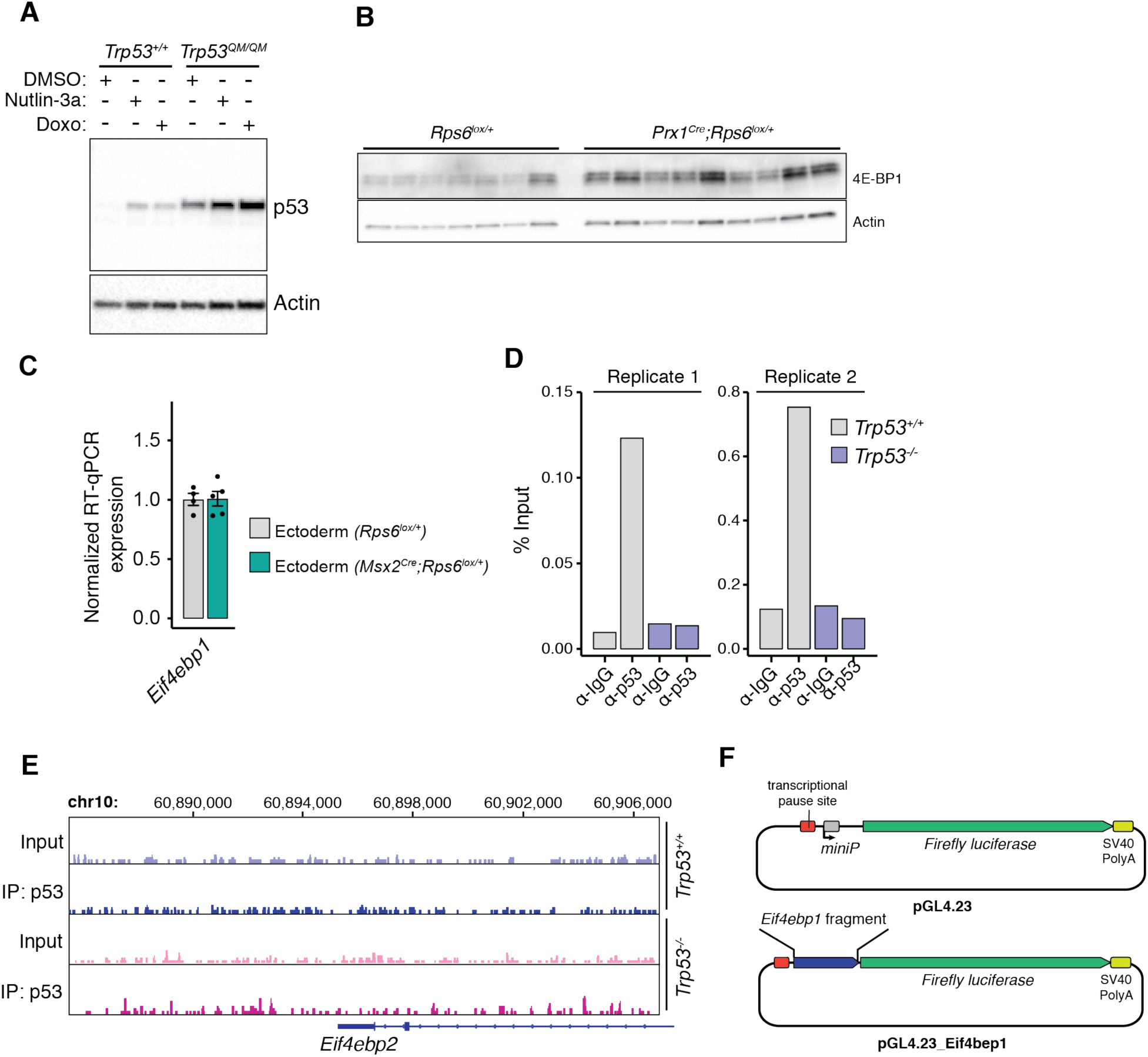
p53 transcriptional activity is required for translational repression, and p53 induces *Eif4ebp1* expression. Related to Figure 5. (**A**) Western blot of p53 stabilization after 8 h of Nutlin-3a or doxorubicin treatment of mouse embryonic fibroblast cells harboring wildtype (*Trp53^+/+^*) or transactivation dead mutant (*Trp53^QM/QM^*) p53. Of note, the *Trp53^QM^* allele also abrogates Mdm2 mediated degradation of p53, thus stabilizing p53 at baseline without drug treatment (Kenzelmann Broz et al., 2013). (**B**) Full Western blot from Figure 5E of 4E-BP1 from E10.5 forelimb mesenchyme cells after removal of the ectoderm layer in *Rps6^lox/+^* and *Prx1^Cre^;Rps6^lox/+^* backgrounds. (**C**) RT-qPCR of *Eif4ebp1* mRNA from ectoderm isolated from E11.5 wildtype (*Rps6^lox/+^*) and *Msx2^Cre^;Rps6^lox/+^* forelimbs. Expression was normalized to *Actb* and *NupL1* and then to wildtype for each tissue. (**D**) ChIP-qPCR of p53 or IgG control from wildtype or *Trp53* null MEFs treated with doxorubicin using gene-specific primers for *Eif4ebp1*. Shown is the percent input of two independent replicates. (**E**) p53 ChIP-Seq gene track of *Eif4ebp2* locus (Kenzelmann Broz et al., 2013). (**F**) Schematic of constructs used to test the p53-dependent promoter activity at the *Eif4ebp1* locus in Figure 5. For all bar plots, error bars = SEM, * *P* < 0.05, ** *P* < 0.01, *** *P* < 0.001, **** *P* < 0.0001, two-tailed t-test, unequal variance.

**Figure S14.**
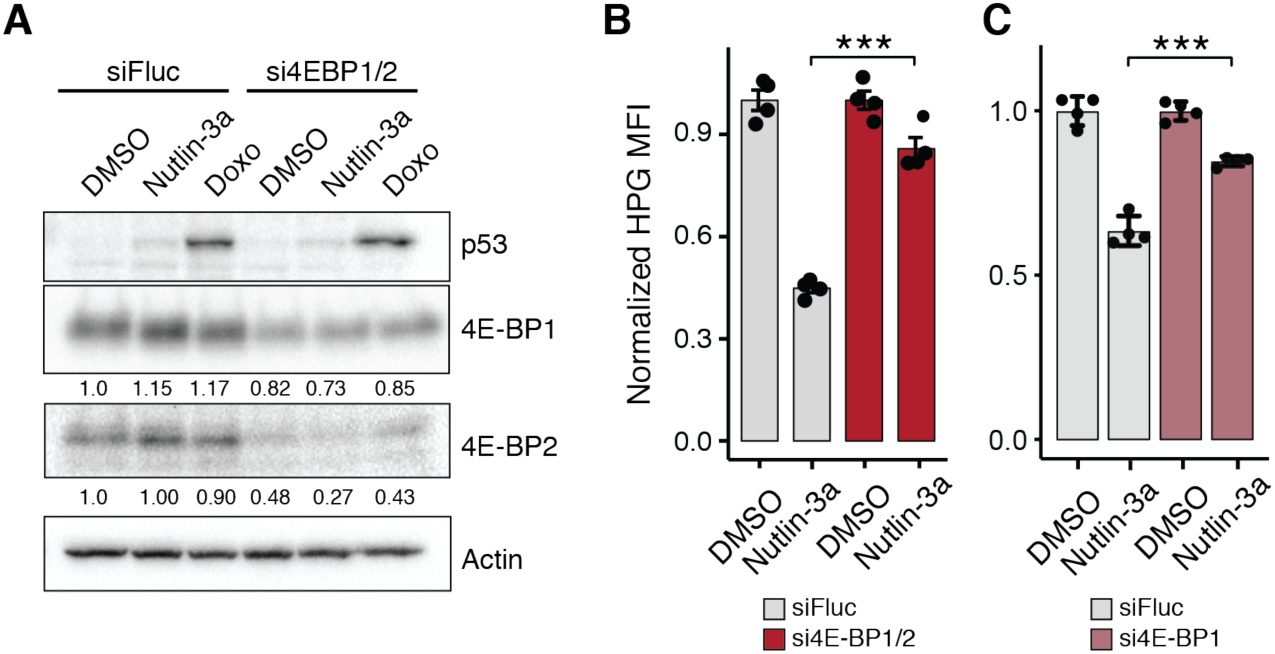
Depletion of 4E-BP1 restores Nutlin-induced translation defects. Related to Figure 5. (**A**) Western blot of 4E-BP1 and 4E-BP2 levels after 16 h of siRNA depletion in NIH3T3 cells. Normalized to DMSO. Numbers indicate the amount of quantified protein relative to siFluc control. (**B**) Quantification of HPG MFI upon combined siRNA knockdown of *Eif4ebp1* and Eif4ebp*2* in NIH3T3 cells normalized to respective DMSO control; *n* = 4. (**C**) Quantification of HPG MFI upon siRNA knockdown of only *Eif4ebp1* in NIH3T3 cells normalized to respective DMSO control; *n* = 4. For all bar plots, error bars = SEM, * *P* < 0.05, ** *P* < 0.01, *** *P* < 0.001, **** *P* < 0.0001, two-tailed t-test, unequal variance.

## REFERENCES

Barlow, J.L., Drynan, L.F., Hewett, D.R., Holmes, L.R., Lorenzo-Abalde, S., Lane, A.L., Jolin, H.E., Pannell, R., Middleton, A.J., Wong, S.H., et al. (2010). A p53-dependent mechanism underlies macrocytic anemia in a mouse model of human 5q-syndrome. Nat. Med. 16, 59–66.

Barna, M., and Niswander, L. (2007). Visualization of cartilage formation: insight into cellular properties of skeletal progenitors and chondrodysplasia syndromes. Dev. Cell 12, 931–941.

Bi, C., Zhang, X., Lu, T., Zhang, X., Wang, X., Meng, B., Zhang, H., Wang, P., Vose, J.M., Chan, W.C., et al. (2017). Inhibition of 4EBP phosphorylation mediates the cytotoxic effect of mechanistic target of rapamycin kinase inhibitors in aggressive B-cell lymphomas. Haematologica 102, 755–764.

Bowen, M.E., McClendon, J., Long, H.K., Sorayya, A., Van Nostrand, J.L., Wysocka, J., and Attardi, L.D. (2019). The Spatiotemporal Pattern and Intensity of p53 Activation Dictates Phenotypic Diversity in p53-Driven Developmental Syndromes. Dev. Cell 50, 212–228.e6.

Brady, C.A., Jiang, D., Mello, S.S., Johnson, T.M., Jarvis, L.A., Kozak, M.M., Kenzelmann Broz, D., Basak, S., Park, E.J., McLaughlin, M.E., et al. (2011). Distinct p53 transcriptional programs dictate acute DNA-damage responses and tumor suppression. Cell 145, 571–583.

Budanov, A.V., and Karin, M. (2008). p53 target genes sestrin1 and sestrin2 connect genotoxic stress and mTOR signaling. Cell 134, 451–460.

Cenik, C., Cenik, E.S., Byeon, G.W., Grubert, F., Candille, S.I., Spacek, D., Alsallakh, B., Tilgner, H., Araya, C.L., Tang, H., et al. (2015). Integrative analysis of RNA, translation, and protein levels reveals distinct regulatory variation across humans. Genome Res. 25, 1610–1621.

Clinton, C., and Gazda, H.T. (1993). Diamond-Blackfan Anemia. In GeneReviews(®), R.A. Pagon, M.P. Adam, H.H. Ardinger, S.E. Wallace, A. Amemiya, L.J. Bean, T.D. Bird, C.-T. Fong, H.C. Mefford, R.J. Smith, et al., eds. (Seattle (WA): University of Washington, Seattle), p.

Dalla Venezia, N., Vincent, A., Marcel, V., Catez, F., and Diaz, J.-J. (2019). Emerging role of eukaryote ribosomes in translational control. Int. J. Mol. Sci. 20.

Danilova, N., Sakamoto, K.M., and Lin, S. (2008). Ribosomal protein S19 deficiency in zebrafish leads to developmental abnormalities and defective erythropoiesis through activation of p53 protein family. Blood 112, 5228–5237.

Deisenroth, C., and Zhang, Y. (2010). Ribosome biogenesis surveillance: probing the ribosomal protein-Mdm2-p53 pathway. Oncogene 29, 4253–4260.

Ding, M., Van der Kwast, T.H., Vellanki, R.N., Foltz, W.D., McKee, T.D., Sonenberg, N., Pandolfi, P.P., Koritzinsky, M., and Wouters, B.G. (2018). The mTOR Targets 4E-BP1/2 Restrain Tumor Growth and Promote Hypoxia Tolerance in PTEN-driven Prostate Cancer. Mol. Cancer Res. 16, 682–695.

Duncan, R., Milburn, S.C., and Hershey, J.W. (1987). Regulated phosphorylation and low abundance of HeLa cell initiation factor eIF-4F suggest a role in translational control. Heat shock effects on eIF-4F. J. Biol. Chem. 262, 380–388.

Dutt, S., Narla, A., Lin, K., Mullally, A., Abayasekara, N., Megerdichian, C., Wilson, F.H., Currie, T., Khanna-Gupta, A., Berliner, N., et al. (2011). Haploinsufficiency for ribosomal protein genes causes selective activation of p53 in human erythroid progenitor cells. Blood 117, 2567–2576.

Flynn, R.A., Martin, L., Spitale, R.C., Do, B.T., Sagan, S.M., Zarnegar, B., Qu, K., Khavari, P.A., Quake, S.R., Sarnow, P., et al. (2015). Dissecting noncoding and pathogen RNA-protein interactomes. RNA 21, 135–143.

Furic, L., Rong, L., Larsson, O., Koumakpayi, I.H., Yoshida, K., Brueschke, A., Petroulakis, E., Robichaud, N., Pollak, M., Gaboury, L.A., et al. (2010). eIF4E phosphorylation promotes tumorigenesis and is associated with prostate cancer progression. Proc Natl Acad Sci USA 107, 14134–14139.

Garami, A., Zwartkruis, F.J.T., Nobukuni, T., Joaquin, M., Roccio, M., Stocker, H., Kozma, S.C., Hafen, E., Bos, J.L., and Thomas, G. (2003). Insulin activation of Rheb, a mediator of mTOR/S6K/4E-BP signaling, is inhibited by TSC1 and 2. Mol. Cell 11, 1457–1466.

Gazda, H.T., Sheen, M.R., Vlachos, A., Choesmel, V., O’Donohue, M.-F., Schneider, H., Darras, N., Hasman, C., Sieff, C.A., Newburger, P.E., et al. (2008). Ribosomal protein L5 and L11 mutations are associated with cleft palate and abnormal thumbs in Diamond-Blackfan anemia patients. Am. J. Hum. Genet. 83, 769–780.

Harfe, B.D., McManus, M.T., Mansfield, J.H., Hornstein, E., and Tabin, C.J. (2005). The RNaseIII enzyme Dicer is required for morphogenesis but not patterning of the vertebrate limb. Proc Natl Acad Sci USA 102, 10898–10903.

Hernandez, O., Way, S., McKenna, J., and Gambello, M.J. (2007). Generation of a conditional disruption of the Tsc2 gene. Genesis 45, 101–106.

Horos, R., Ijspeert, H., Pospisilova, D., Sendtner, R., Andrieu-Soler, C., Taskesen, E., Nieradka, A., Cmejla, R., Sendtner, M., Touw, I.P., et al. (2012). Ribosomal deficiencies in Diamond-Blackfan anemia impair translation of transcripts essential for differentiation of murine and human erythroblasts. Blood 119, 262–272.

Hurst, J.A., Baraitser, M., and Wonke, B. (1991). Autosomal dominant transmission of congenital erythroid hypoplastic anemia with radial abnormalities. Am. J. Med. Genet. 40, 482–484.

Ingolia, N.T., Lareau, L.F., and Weissman, J.S. (2011). Ribosome profiling of mouse embryonic stem cells reveals the complexity and dynamics of mammalian proteomes. Cell 147, 789–802.

Ingolia, N.T., Brar, G.A., Rouskin, S., McGeachy, A.M., and Weissman, J.S. (2012). The ribosome profiling strategy for monitoring translation in vivo by deep sequencing of ribosome-protected mRNA fragments. Nat. Protoc. 7, 1534–1550.

Jaako, P., Debnath, S., Olsson, K., Bryder, D., Flygare, J., and Karlsson, S. (2012). Dietary L-leucine improves the anemia in a mouse model for Diamond-Blackfan anemia. Blood 120, 2225–2228.

Jacks, T., Remington, L., Williams, B.O., Schmitt, E.M., Halachmi, S., Bronson, R.T., and Weinberg, R.A. (1994). Tumor spectrum analysis in p53-mutant mice. Curr. Biol. 4, 1–7.

Jones, N.C., Lynn, M.L., Gaudenz, K., Sakai, D., Aoto, K., Rey, J.-P., Glynn, E.F., Ellington, L., Du, C., Dixon, J., et al. (2008). Prevention of the neurocristopathy Treacher Collins syndrome through inhibition of p53 function. Nat. Med. 14, 125–133.

Kastenhuber, E.R., and Lowe, S.W. (2017). Putting p53 in context. Cell 170, 1062–1078.

Kasteri, J., Das, D., Zhong, X., Persaud, L., Francis, A., Muharam, H., and Sauane, M. (2018). Translation Control by p53. Cancers (Basel) 10.

Kenzelmann Broz, D., Spano Mello, S., Bieging, K.T., Jiang, D., Dusek, R.L., Brady, C.A., Sidow, A., and Attardi, L.D. (2013). Global genomic profiling reveals an extensive p53-regulated autophagy program contributing to key p53 responses. Genes Dev. 27, 1016–1031.

Khajuria, R.K., Munschauer, M., Ulirsch, J.C., Fiorini, C., Ludwig, L.S., McFarland, S.K., Abdulhay, N.J.,Specht, H., Keshishian, H., Mani, D.R., et al. (2018). Ribosome levels selectively regulate translation and lineage commitment in human hematopoiesis. Cell 173, 90–103.e19.

Kongsuwan, K., Yu, Q., Vincent, A., Frisardi, M.C., Rosbash, M., Lengyel, J.A., and Merriam, J. (1985). A Drosophila Minute gene encodes a ribosomal protein. Nature 317, 555–558.

Langmead, B., and Salzberg, S.L. (2012). Fast gapped-read alignment with Bowtie 2. Nat. Methods 9, 357–359.

Law, C.W., Chen, Y., Shi, W., and Smyth, G.K. (2014). voom: Precision weights unlock linear model analysis tools for RNA-seq read counts. Genome Biol. 15, R29.

Lee, B., Vissing, H., Ramirez, F., Rogers, D., and Rimoin, D. (1989). Identification of the molecular defect in a family with spondyloepiphyseal dysplasia. Science 244, 978–980.

Lee, C.-H., Kiparaki, M., Blanco, J., Folgado, V., Ji, Z., Kumar, A., Rimesso, G., and Baker, N.E. (2018). A regulatory response to ribosomal protein mutations controls translation, growth, and cell competition. Dev. Cell 46, 456–469.e4.

Lin, T.A., Kong, X., Haystead, T.A., Pause, A., Belsham, G., Sonenberg, N., and Lawrence, J.C. (1994). PHAS-I as a link between mitogen-activated protein kinase and translation initiation. Science 266, 653– 656.

Loayza-Puch, F., Drost, J., Rooijers, K., Lopes, R., Elkon, R., and Agami, R. (2013). p53 induces transcriptional and translational programs to suppress cell proliferation and growth. Genome Biol. 14, R32.

Logan, M., Martin, J.F., Nagy, A., Lobe, C., Olson, E.N., and Tabin, C.J. (2002). Expression of Cre Recombinase in the developing mouse limb bud driven by a Prxl enhancer. Genesis 33, 77–80.

Ludwig, L.S., Gazda, H.T., Eng, J.C., Eichhorn, S.W., Thiru, P., Ghazvinian, R., George, T.I., Gotlib, J.R., Beggs, A.H., Sieff, C.A., et al. (2014). Altered translation of GATA1 in Diamond-Blackfan anemia. Nat. Med. 20, 748–753.

Lufkin, T. (2007). In situ hybridization of whole-mount mouse embryos with RNA probes: hybridization, washes, and histochemistry. CSH Protoc. 2007, pdb.prot4823.

Mamane, Y., Petroulakis, E., Rong, L., Yoshida, K., Ler, L.W., and Sonenberg, N. (2004). eIF4E--from translation to transformation. Oncogene 23, 3172–3179.

Marine, J.-C.W., Dyer, M.A., and Jochemsen, A.G. (2007). MDMX: from bench to bedside. J. Cell Sci. 120, 371–378.

Martin, M. (2011). Cutadapt removes adapter sequences from high-throughput sequencing reads. EMBnet j. 17, 10.

McGlincy, N.J., and Ingolia, N.T. (2017). Transcriptome-wide measurement of translation by ribosome profiling. Methods 126, 112–129.

McGowan, K.A., and Mason, P.J. (2011). Animal models of Diamond Blackfan anemia. Semin. Hematol. 48, 106–116.

McGowan, K.A., Li, J.Z., Park, C.Y., Beaudry, V., Tabor, H.K., Sabnis, A.J., Zhang, W., Fuchs, H., de Angelis, M.H., Myers, R.M., et al. (2008). Ribosomal mutations cause p53-mediated dark skin and pleiotropic effects. Nat. Genet. 40, 963–970.

McGowan, K.A., Pang, W.W., Bhardwaj, R., Perez, M.G., Pluvinage, J.V., Glader, B.E., Malek, R., Mendrysa, S.M., Weissman, I.L., Park, C.Y., et al. (2011). Reduced ribosomal protein gene dosage and p53 activation in low-risk myelodysplastic syndrome. Blood 118, 3622–3633.

Merico, D., Isserlin, R., Stueker, O., Emili, A., and Bader, G.D. (2010). Enrichment map: a network-based method for gene-set enrichment visualization and interpretation. PLoS ONE 5, e13984.

Morgado-Palacin, L., Varetti, G., Llanos, S., Gómez-López, G., Martinez, D., and Serrano, M. (2015). Partial Loss of Rpl11 in Adult Mice Recapitulates Diamond-Blackfan Anemia and Promotes Lymphomagenesis. Cell Rep. 13, 712–722.

Mo, R., Freer, A.M., Zinyk, D.L., Crackower, M.A., Michaud, J., Heng, H.H., Chik, K.W., Shi, X.M., Tsui, L.C., Cheng, S.H., et al. (1997). Specific and redundant functions of Gli2 and Gli3 zinc finger genes in skeletal patterning and development. Development 124, 113–123.

Muzumdar, M.D., Tasic, B., Miyamichi, K., Li, L., and Luo, L. (2007). A global double-fluorescent Cre reporter mouse. Genesis 45, 593–605.

Narla, A., and Ebert, B.L. (2010). Ribosomopathies: human disorders of ribosome dysfunction. Blood 115, 3196–3205.

Panić, L., Tamarut, S., Sticker-Jantscheff, M., Barkić, M., Solter, D., Uzelac, M., Grabusić, K., and Volarević, S. (2006). Ribosomal protein S6 gene haploinsufficiency is associated with activation of a p53-dependent checkpoint during gastrulation. Mol. Cell. Biol. 26, 8880–8891.

Payne, E.M., Virgilio, M., Narla, A., Sun, H., Levine, M., Paw, B.H., Berliner, N., Look, A.T., Ebert, B.L., and Khanna-Gupta, A. (2012). L-Leucine improves the anemia and developmental defects associated with Diamond-Blackfan anemia and del(5q) MDS by activating the mTOR pathway. Blood 120, 2214– 2224.

Pelletier, J., Graff, J., Ruggero, D., and Sonenberg, N. (2015). Targeting the eIF4F translation initiation complex: a critical nexus for cancer development. Cancer Res. 75, 250–263.

Ritchie, M.E., Phipson, B., Wu, D., Hu, Y., Law, C.W., Shi, W., and Smyth, G.K. (2015). limma powers differential expression analyses for RNA-sequencing and microarray studies. Nucleic Acids Res. 43, e47.

Robinson, M.D., McCarthy, D.J., and Smyth, G.K. (2010). edgeR: a Bioconductor package for differential expression analysis of digital gene expression data. Bioinformatics 26, 139–140.

Ruggero, D., Montanaro, L., Ma, L., Xu, W., Londei, P., Cordon-Cardo, C., and Pandolfi, P.P. (2004). The translation factor eIF-4E promotes tumor formation and cooperates with c-Myc in lymphomagenesis. Nat. Med. 10, 484–486.

Saxton, R.A., and Sabatini, D.M. (2017). mTOR Signaling in Growth, Metabolism, and Disease. Cell 168, 960–976.

Sekiyama, N., Arthanari, H., Papadopoulos, E., Rodriguez-Mias, R.A., Wagner, G., and Léger-Abraham, M. (2015). Molecular mechanism of the dual activity of 4EGI-1: Dissociating eIF4G from eIF4E but stabilizing the binding of unphosphorylated 4E-BP1. Proc Natl Acad Sci USA 112, E4036–45.

Signer, R.A.J., Magee, J.A., Salic, A., and Morrison, S.J. (2014). Haematopoietic stem cells require a highly regulated protein synthesis rate. Nature 509, 49–54.

Smith, T., Heger, A., and Sudbery, I. (2017). UMI-tools: modeling sequencing errors in Unique Molecular Identifiers to improve quantification accuracy. Genome Res. 27, 491–499.

Sonenberg, N., and Hinnebusch, A.G. (2009). Regulation of translation initiation in eukaryotes: mechanisms and biological targets. Cell 136, 731–745.

Sulic, S., Panic, L., Barkic, M., Mercep, M., Uzelac, M., and Volarevic, S. (2005). Inactivation of S6 ribosomal protein gene in T lymphocytes activates a p53-dependent checkpoint response. Genes Dev. 19, 3070–3082.

Sun, X., Lewandoski, M., Meyers, E.N., Liu, Y.H., Maxson, R.E., and Martin, G.R. (2000). Conditional inactivation of Fgf4 reveals complexity of signalling during limb bud development. Nat. Genet. 25, 83–86.

Thoreen, C.C., Chantranupong, L., Keys, H.R., Wang, T., Gray, N.S., and Sabatini, D.M. (2012). A unifying model for mTORC1-mediated regulation of mRNA translation. Nature 485, 109–113.

Truitt, M.L., Conn, C.S., Shi, Z., Pang, X., Tokuyasu, T., Coady, A.M., Seo, Y., Barna, M., and Ruggero, D. (2015). Differential Requirements for eIF4E Dose in Normal Development and Cancer. Cell 162, 59–71.

Twigg, S.R.F., Lloyd, D., Jenkins, D., Elçioglu, N.E., Cooper, C.D.O., Al-Sannaa, N., Annagür, A., Gillessen-Kaesbach, G., Hüning, I., Knight, S.J.L., et al. (2012). Mutations in multidomain protein MEGF8 identify a Carpenter syndrome subtype associated with defective lateralization. Am. J. Hum. Genet. 91, 897–905.

Vassilev, L.T., Vu, B.T., Graves, B., Carvajal, D., Podlaski, F., Filipovic, Z., Kong, N., Kammlott, U., Lukacs, C., Klein, C., et al. (2004). In vivo activation of the p53 pathway by small-molecule antagonists of MDM2. Science 303, 844–848.

Vlachos, A., Rosenberg, P.S., Atsidaftos, E., Alter, B.P., and Lipton, J.M. (2012). Incidence of neoplasia in Diamond Blackfan anemia: a report from the Diamond Blackfan Anemia Registry. Blood 119, 3815– 3819.

Volarevic, S., Stewart, M.J., Ledermann, B., Zilberman, F., Terracciano, L., Montini, E., Grompe, M., Kozma, S.C., and Thomas, G. (2000). Proliferation, but not growth, blocked by conditional deletion of 40S ribosomal protein S6. Science 288, 2045–2047.

Wang, S., Konorev, E.A., Kotamraju, S., Joseph, J., Kalivendi, S., and Kalyanaraman, B. (2004). Doxorubicin induces apoptosis in normal and tumor cells via distinctly different mechanisms. intermediacy of H(2)O(2)- and p53-dependent pathways. J. Biol. Chem. 279, 25535–25543.

Weinberg, D.E., Shah, P., Eichhorn, S.W., Hussmann, J.A., Plotkin, J.B., and Bartel, D.P. (2016). Improved Ribosome-Footprint and mRNA Measurements Provide Insights into Dynamics and Regulation of Yeast Translation. Cell Rep. 14, 1787–1799.

Wu, D., and Smyth, G.K. (2012). Camera: a competitive gene set test accounting for inter-gene correlation. Nucleic Acids Res. 40, e133.

Yelick, P.C., and Trainor, P.A. (2015). Ribosomopathies: Global process, tissue specific defects. Rare Dis. 3, e1025185.

Zeller, R., López-Ríos, J., and Zuniga, A. (2009). Vertebrate limb bud development: moving towards integrative analysis of organogenesis. Nat. Rev. Genet. 10, 845–858.

Zhang, Y., Nicholatos, J., Dreier, J.R., Ricoult, S.J.H., Widenmaier, S.B., Hotamisligil, G.S., Kwiatkowski, D.J., and Manning, B.D. (2014). Coordinated regulation of protein synthesis and degradation by mTORC1. Nature 513, 440–443.

